# IntegrateRigor: annotation-free integration optimization for cell identity recovery reveals cancer–immune interface niches

**DOI:** 10.64898/2026.05.14.725078

**Authors:** Zhiqian Zhai, Changhu Wang, Chengfeng Jiang, Ziqi Rong, Jingyi Jessica Li

**Affiliations:** Department of Statistics and Data Science, University of California, Los Angeles, CA 90095; Biostatistics Program, Public Health Science Division, Fred Hutchinson Cancer, Seattle, WA 98109; Paul G. Allen School of Computer Science and Engineering, University of Washington, Seattle, WA 98195; Department of Biostatistics, University of Washington, Seattle, WA 98195

## Abstract

Integrating single-cell and spatial transcriptomics data across batches is essential for recovering comparable cell identities—including cell types, subtypes, and states—as a prerequisite for downstream analyses in multi-condition and large-scale studies. This task remains challenging because between-batch variation removal often conflicts with cell identity preservation, and current methods typically rely on generic highly variable gene selection and lack principled metrics for hyperparameter tuning when cell identity annotations are unavailable. Together, these limitations often lead to over-integration, which merges biologically distinct cell identities, or under-integration, which leaves cells separated by batch rather than identity. Here we introduce IntegrateRigor, a data-driven, annotation-free, method-agnostic framework that optimizes integration specifically for reliable cell identity recovery across batches. IntegrateRigor first selects genes whose expression patterns are stable across batches using a gene-wise likelihood-based batch stability score, excluding batch-sensitive genes that can bias cell identity alignment during integration. It then identifies the optimal integration configuration across methods and hyperparameters by defining a dataset-level integration score that explicitly balances between-batch variation removal against cell identity preservation, without requiring prior annotations. In a colorectal cancer single-cell and spatial transcriptomics dataset, IntegrateRigor revealed previously uncharacterized cancer–immune interface niches in the tumor microenvironment that were masked by under-integration under default settings and by over-integration in previous literature. Across diverse datasets spanning multiple sources of between-batch variation, IntegrateRigor consistently improved cell identity recovery by mitigating both over-integration and under-integration across five state-of-the-art integration methods. By transforming integration from a heuristic preprocessing step into a statistically principled, dataset-adaptive procedure for cell identity recovery, IntegrateRigor improves the reproducibility and biological discovery power of large-scale single-cell and spatial transcriptomics analyses.

## Introduction

Large-scale single-cell RNA sequencing (scRNA-seq) and spatial transcriptomics (ST) are transforming biology by enabling systematic measurements of gene expression at single-cell and spatial resolution [1–3]. The increasing availability of large reference atlases [4–6], together with the rapid accumulation of datasets generated across diverse experimental and biological sources, has greatly expanded opportunities to study cellular heterogeneity, tissue organization, and disease processes at scale. Realizing this potential, however, increasingly requires integrating data across multiple batches to recover comparable cell identities, a step that remains technically and conceptually challenging.

Here, a *batch* refers broadly to a sample or group of cells collected as a replicate or under a distinct condition, such as from different donors, time points, laboratories, sequencing platforms, or even species [7]. Throughout this manuscript, *cell identity* denotes the transcriptomic characteristics of a cell, including its type, subtype, or state [8]. From a computational perspective, consistent with recent work [9, 10], cell identities are manifested as high-density regions in the distribution of cells in a latent embedding space. *Between-batch differences* can arise from two distinct sources: (1) differences in cell identity composition across batches, and (2) differences in measured gene expression for the same cell identity across batches. The former does not hinder cell identity recovery, as cells with the same identity remain comparable in their expression profiles across batches. The latter, which we term *within-identity between-batch expression variation* and hereafter abbreviate as *between-batch variation*, obscures shared cell identity structure and is the primary challenge for data integration. Importantly, between-batch variation is not limited to technical artifacts but can also reflect biological differences between batches, such as cell-type-specific differential expression. Simply pooling cells across batches before identifying cell identities is not a valid option, as it fails to account for between-batch variation and thereby compromises cell identity recovery, as well as all downstream analyses that depend on it, such as differential expression testing, trajectory and lineage inference, cell–cell communication analysis, and cell-identity-specific eQTL [11–14].

Integration is therefore essential for constructing a shared space in which cell identities can be consistently recovered and compared across batches [15, 16]. Without proper integration, cells with the same identity but from different batches may remain separated due to between-batch variation [17, 18]. Crucially, cell identity recovery is a prerequisite for any cell-identity–based downstream analysis; these analyses should be performed only after reliable cell identities have been established across batches, not entangled with the integration step itself. Effective integration must therefore balance two competing objectives: preserving cell identity structure while minimizing between-batch variation. Failure to achieve this balance leads to two opposite failure modes. *Under-integration* leaves residual between-batch variation, causing cells to group by batch rather than by true identity. *Over-integration*, in contrast, removes biologically meaningful between-cell-identity variation and artificially merges distinct cell identities [7, 19–21]. Both distort cell identity recovery and undermine the rigor and reproducibility of any downstream analyses. A wide range of computational methods has been developed for single-cell data integration, including Harmony [22], Seurat-RPCA [23], scVI [24], FastMNN [25], and LIGER [26], among many others [12, 27–29]. These methods are primarily designed to align cells with the same biological identity across batches into a shared low-dimensional embedding. Other approaches, such as Crescendo [12] and CellANOVA [30], take the integrated embedding as input to perform count-level batch correction and cell-type-specific differential expression analysis, respectively; their outputs therefore depend directly on the quality of the integrated embedding. However, despite substantial methodological progress, current integration practice remains limited in two key respects, especially in the common setting where accurate cell identity annotations are unavailable before integration [7].

First, the choice of input genes is a key but underappreciated determinant of integration quality for cell identity recovery [7, 31]. Most integration methods begin with generic gene selection, typically using highly variable genes (HVGs) selected either within each batch and then pooled, or from cells pooled across batches [31, 32]. However, these HVGs may be sensitive to between-batch variation; that is, their expression levels for the same cell identity may differ substantially across batches. For example, many HVGs primarily reflect activity or stress-response programs rather than intrinsic cell identity [33, 34], and are therefore highly sensitive to between-batch variation (for example, mitochondrial and ribosomal genes) [31, 35]. Using such HVGs introduces between-batch variation into the integration input, which hinders reliable cell identity alignment across batches. In contrast, genes tied to conserved cell-identity programs show relatively stable expression patterns across donors, tissues, and even species, providing a biologically grounded pool of candidate genes [36–39]. A recent benchmarking study [31] showed that gene selection can be as consequential as method choice for integration quality. Although a cell-type-annotation-dependent metric, group technical effects (GTE), has been proposed to quantify genewise between-batch variation [40], reliable annotations are often unavailable before integration, and using them for gene selection introduces circularity into the workflow.

Second, existing metrics for evaluating the trade-off between batch mixing and cell identity preservation fall into two categories, each with important limitations. First, annotation-dependent metrics—such as cell-type local inverse Simpson’s index (cLISI) [17, 41] and adjusted Rand index (ARI) [41, 42] for cell identity preservation, and cell-type-aware integration LISI (CiLISI) [17] for batch mixing—are informative when accurate cell identity labels are available. However, they create a circular dependency: integration is performed precisely to recover cell identities across batches, yet these metrics presuppose that reliable annotations already exist. Second, annotation-free metrics—such as integration LISI (iLISI), k-nearest-neighbor batch-effect test (kBET), principal component regression (PCR), and batch average silhouette width (batch ASW) [17, 21, 22, 41]—avoid this dependency but measure only batch mixing without assessing cell identity preservation, and are therefore unable to detect over-integration. Although the silhouette score is an annotation-free metric that can assess cell cluster separation and thus cell identity preservation, it depends on the performance of the clustering algorithm and has been shown to be unreliable in single-cell integration benchmarking [41]. As a result, no existing annotation-free approach jointly assesses batch mixing and cell identity preservation to guide integration decisions for a specific dataset. In practice, researchers often resort to subjective visual inspection of UMAP plots [43] and trial-and-error hyperparameter tuning (e.g., the diversity clustering penalty in Harmony), making integration difficult to reproduce and to optimize for a given dataset.

Together, these limitations reveal a critical gap: no existing approach jointly addresses gene selection, hyperparameter optimization, and method selection in a statistically principled, annotation-free, dataset-specific manner with the explicit goal of reliable cell identity recovery. To address this gap, we propose IntegrateRigor, a data-driven, method-agnostic framework for optimizing single-cell and spatial transcriptomics integration for cell identity recovery. IntegrateRigor is not a standalone integration method; rather, it provides a principled wrapper for evaluating and tuning existing integration methods. It first identifies *batch-stable genes (BSGs)*—genes whose expression patterns across cell identities remain consistent across batches—and selects HVGs within this batch-stable set as cell-identity-informative input features for integration. Gene selection is achieved through a likelihood-based, annotation-free per-gene *batch stability score (BSS)*, which quantifies gene-wise stability across batches without requiring cell identity annotations. After integration, IntegrateRigor evaluates the resulting cell embedding using two annotation-free metrics: the *batch alignment score (BAS)*, which measures residual between-batch variation, and the *cell identity score (CIS)*, which measures preservation of cell identity structure. These two scores are aggregated into a dataset-level *integration score* that identifies the optimal configuration (i.e., method and hyperparameters) for reliable cell identity recovery on a given dataset. This shifts the focus from global method benchmarking to dataset-specific optimization: although benchmarking studies provide valuable overall rankings of integration methods [17, 18, 35], they typically use fixed parameter settings across datasets and therefore may not reflect each method’s best achievable performance on any given dataset. By jointly addressing gene selection, hyperparameter tuning, and method selection, IntegrateRigor transforms integration from a heuristic, trial-and-error workflow into a rigorous, annotation-free, and data-adaptive procedure for cell identity recovery.

In an application to colorectal cancer single-cell and spatial transcriptomics [12], IntegrateRigor enabled reliable cell identity recovery that revealed previously unannotated cancer–immune interface niches in the tumor microenvironment, structure that was obscured under both default integration settings and the over-integrated embedding reported in previous literature. Across six integration tasks spanning diverse sources of between-batch variation (i.e., sequencing platforms, sample conditions, developmental stage, species, and spatial tissue sections), IntegrateRigor consistently improved cell identity recovery of five widely used methods (Seurat-RPCA, Harmony, scVI, FastMNN, and LIGER). Notably, Harmony, which showed only moderate performance under default settings, achieved one of the best overall results when paired with IntegrateRigor, demonstrating that dataset-specific optimization can unlock the full potential of existing methods for cell identity recovery.

## Results

### Overview of IntegrateRigor: an annotation-free framework for batch-stable gene selection and integration optimization for cell identity recovery

IntegrateRigor provides an annotation-free, data-adaptive wrapper that optimizes data integration for reliable cell identity recovery while minimizing human subjectivity in hyperparameter tuning. By combining BSG selection with data-driven hyperparameter optimization, IntegrateRigor improves cell identity recovery by mitigating both under-integration and over-integration, without requiring prior cell identity annotations. This design makes IntegrateRigor broadly applicable across datasets and easy to use as a plug-in within existing integration pipelines. Details of the IntegrateRigor method are described in the **Methods Section**.

As illustrated in Figure 1, IntegrateRigor introduces annotation-free statistical metrics at three levels: (1) a likelihood-based per-gene BSS that selects batch-stable, cell-identity-informative genes as input features; (2) likelihood-based embedding-level metrics, BAS and CIS, that jointly quantify batch mixing and cell identity preservation; and (3) a dataset-level integration score that identifies the optimal method and hyperparameter setting for a given dataset.

**Figure 1:**
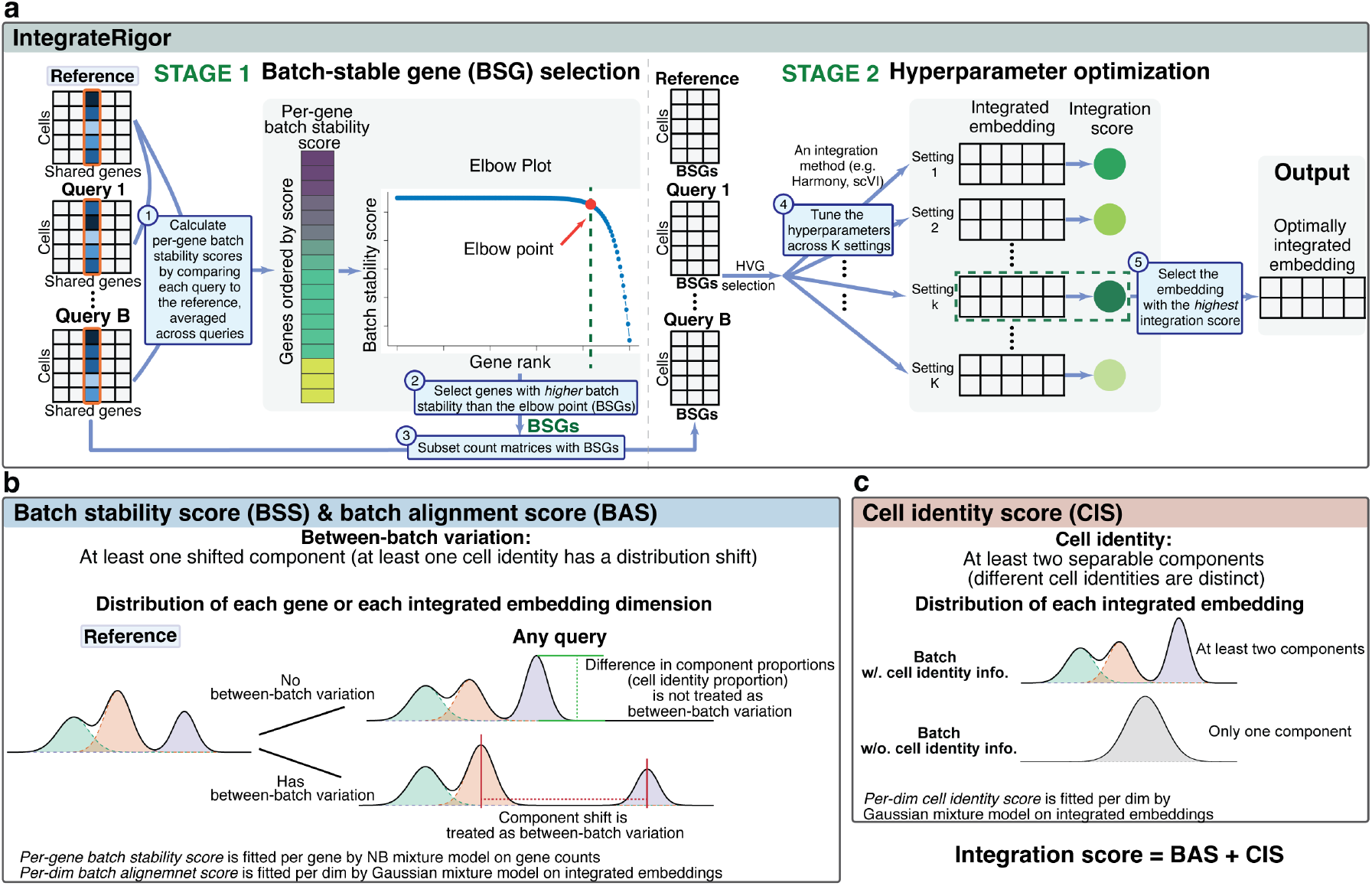
Overview of IntegrateRigor framework. **a**. Schematic of the IntegrateRigor workflow. In Stage 1, batch-stable genes (BSGs) are selected by comparing each query batch to a reference batch and computing a per-gene batch stability score (BSS), which quantifies the extent to which gene-level distributions are preserved across batches. Genes ranked above the elbow point are retained as BSGs. In Stage 2, count matrices are restricted to BSGs, and candidate hyperparameter settings for a chosen integration method are evaluated. The integrated embedding with the highest integration score is selected as optimal. **b**. Illustration of the BSS and batch alignment score (BAS). For each gene or each integrated embedding dimension, between-batch variation is modeled as a distributional shift in at least one mixture component corresponding to a shared cell identity across batches. Differences in mixture component proportions are not treated as between-batch variation. BSS is estimated on gene counts using a negative binomial mixture model, whereas per-dimension BASs are estimated on integrated embeddings using a Gaussian mixture model. **c**. Illustration of the cell identity score (CIS). For each integrated embedding dimension, preservation of cell identity is quantified by the presence of multiple Gaussian mixture components, indicating distinguishable cell identities. The per-dimension CIS is estimated using a Gaussian mixture model on the integrated embedding. The overall integration score is defined as the sum of the median BAS and median CIS across embedding dimensions.

In Stage 1, IntegrateRigor identifies a set of BSGs to use as input features for downstream integration (Figure 1a). Here, between-batch variation refers to differences in measured gene expression for the same cell identity across batches, as distinct from differences in cell identity composition. We assume that a subset of genes carries cell identity information while showing minimal between-batch variation; that is, their expression within each cell identity does not shift substantially across batches (e.g., they are not cell-identity-specific differentially expressed genes across batches), and IntegrateRigor aims to identify and select these genes. Intuitively, such a gene should show the same relative expression patterns across cell identities in every batch, even if the proportions of cell identities differ between batches.

To formalize this intuition, we construct a likelihood-based statistic to quantify the degree to which a gene’s expression distribution is preserved across batches. The idea is to leverage a *reference batch*—either user-defined (e.g., an atlas or a well-annotated anchor dataset) or, if not specified, automatically selected as the batch with the greatest cell-identity diversity, approximated by the largest number of cell clusters within a batch—to guide recovery of shared cell identity structure across batches. We refer to the remaining batches as *query batches*. Specifically, for each gene and each query batch, we fit two mixture models: a reference-constrained model that fixes the mixture components (corresponding to cell identities) as estimated in the reference batch while allowing only component proportions to differ, and an unconstrained model in which both components and proportions are free to vary from the reference batch. The difference in model fit between these two models, measured by the normalized log-likelihood, quantifies the degree of between-batch variation for that gene in that query batch; averaging this difference across all query batches yields the per-gene BSS, which captures how consistently a gene’s expression pattern is maintained across batches (Figure 1b). Genes with higher scores are selected as BSGs using a data-adaptive threshold determined by elbow-point detection, yielding input features that preferentially retain cell identity signals while reducing between-batch variation. HVGs are then selected within the BSGs as final input features, ensuring that they are both cell-identity-informative and stable across batches.

In Stage 2, IntegrateRigor identifies the optimal integration result among candidate embeddings generated under different hyperparameter settings (Figure 1a). Each embedding is evaluated using two complementary annotation-free metrics that correspond to the two objectives of integration for cell identity recovery. We first define BAS and CIS for each dimension of the integrated embedding. The BAS quantifies the degree of between-batch alignment in each embedding dimension, where higher values reflect better batch mixing. The CIS measures how strongly each embedding dimension preserves biologically meaningful cell identity structure, quantified by the degree to which the distribution of embedding values in that dimension reflects distinct cell subpopulations rather than a unimodal background. Specifically, for each embedding dimension, we compute the log-likelihood difference between a constrained and an unconstrained model separately for the reference batch and each query batch, then average across batches, with each per-batch score normalized by cell number to ensure comparability. We then aggregate the per-dimension BAS and CIS into dataset-level scores by taking the median across embedding dimensions. This aggregation is justified by the near-orthogonality of the embedding dimensions: median absolute Pearson correlations across dimension pairs ranged from 0.025 to 0.068 across integration methods (Figure S7), indicating that the dimensions contribute largely independent information. Finally, the dataset-level integration score is defined as the sum of dataset-level BAS and CIS, directly capturing the trade-off between batch mixing and cell identity preservation: BAS penalizes under-integration, whereas CIS penalizes over-integration (Figure 1b–c). Because both scores are defined as normalized log-likelihood differences, they are on a comparable scale and can be summed directly, without requiring an additional weighting parameter. The hyperparameter setting that maximizes the integration score is then selected as the optimal configuration for cell identity recovery on the given dataset.

Empirical validation of the statistical models underlying IntegrateRigor is in the **Methods Section**.

### Batch stability scores identify batch-stable genes and improve cell identity recovery by excluding genes confounded by sample and technical variation

The BSS, computed in Stage 1 of IntegrateRigor, quantifies how stably a gene’s expression pattern is maintained across batches within the same cell identity. Genes with high BSS show consistent expression patterns across batches within the same cell identity; genes with low BSS show substantial cross-batch shifts, indicating that their expression is strongly influenced by between-batch variation and may obscure shared cell identity structure, thereby hindering reliable cell identity recovery. We validated the BSS and evaluated the downstream benefit of BSS-based gene selection using the intestinal failure-associated liver disease (IFALD) dataset comprising three donors [44].

Genes ranked highly by the BSS showed more concordant expression patterns across batches, whereas low-ranked genes displayed substantial between-batch variation (Figure 2a). Principal component analysis of the pooled cells confirmed this separation: genes with large loadings on the batch-dominated principal component 1 (PC1) tended to have lower BSS and were not classified as batch-stable, whereas genes with large loadings on the cell-type-dominated PC2 tended to have higher BSS and were classified as batch-stable (Figure S9). To benchmark the BSS against an existing annotation-dependent metric, we compared the BSS with the GTE score [40], which requires cell-type annotations. BSS was linearly correlated with GTE on the log scale across all datasets (Pearson *r* = 0.50–0.69; all *p* < 2.2 × 10^−16^; Figure S10), confirming that BSS captures a similar notion of between-batch variation as this annotation-dependent metric, while requiring no cell-type labels.

**Figure 2:**
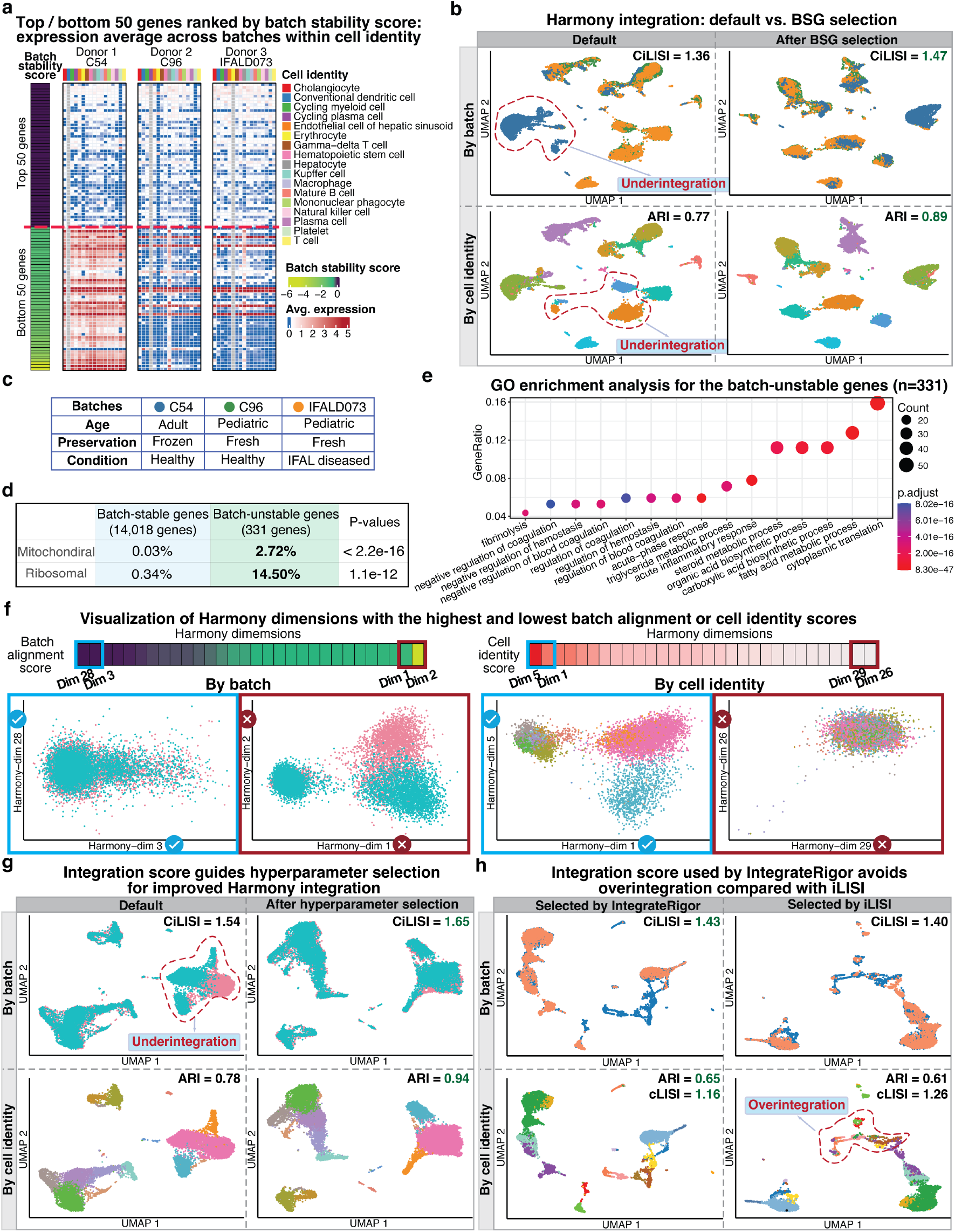
IntegrateRigor identifies batch-unstable genes and uses integration scores to improve gene selection and hyperparameter selection while avoiding overintegration. **a**. Heatmaps of the top and bottom 50 genes ranked by BSS, showing average expression across batches within each cell identity. Top-ranked genes are batch-stable, whereas bottom-ranked genes exhibit substantial batch-dependent variability. **b**. Harmony integration before and after selecting highly variable genes from the BSGs set. BSG selection improves batch mixing (higher CiLISI) while better preserving cell identity structure (higher ARI), reducing underintegration. **c**. Summary of the three IFALD batches, which differ in donors’ age, conditions, and sample preservation. **d**. Comparison of mitochondrial and ribosomal gene proportions between batch-stable and batch-unstable genes, showing strong enrichment of ribosomal genes among the batch-unstable genes. Here, the batch-unstable genes are the genes with low BSS scores and are excluded from the integration. **e**. Gene Ontology enrichment analysis of the batch-unstable genes, highlighting biological processes related to coagulation, inflammatory response, lipid metabolism, and cytoplasmic translation. **f**. Visualization of Harmony dimensions with the highest and lowest batch alignment or CISs. Dimensions with high BASs show better batch mixing, whereas dimensions with high CISs better preserve biological structure. **g**. Integration scores guide hyperparameter selection for Harmony, improving both batch correction and cell identity preservation relative to the default setting. **h**. Compared with selection based on iLISI, IntegrateRigor selects a solution that avoids overintegration and better preserves cell identity, as reflected by a lower cLISI. Cell identity labels and color annotations for the UMAPs are provided in Supplementary Figures S13–S15.

IntegrateRigor ranks genes by BSS and uses a data-adaptive elbow-point threshold [45] to exclude genes that are clearly not batch-stable, retaining the remainder as BSGs for downstream HVG selection. This approach is deliberately conservative: rather than aggressively selecting only the most stable genes, it removes the clearly problematic ones while preserving a broad pool for HVG selection. On the IFALD dataset, only 331 of 14,349 genes were excluded as batch-unstable, leaving 14,018 BSGs. In both the default and BSG-restricted settings, the same number of 2,000 HVGs were used as input features; the only difference was whether those HVGs were drawn from the full gene set or restricted to BSGs. When Harmony was applied to the default 2,000 HVGs, the integrated embedding showed clear under-integration, with cells of the same type remaining separated by batch. Restricting HVG selection to BSGs substantially improved cell identity recovery, increasing CiLISI from 1.33 to 1.47 and ARI from 0.77 to 0.89 (Figure 2b). Consistent improvements were observed across all five integration methods evaluated (FastMNN, Harmony, LIGER, scVI, and Seurat-RPCA), demonstrating that BSG selection is a modular, method-agnostic improvement that can be plugged into existing pipelines.

To understand why batch-unstable genes hinder cell identity recovery, we examined their biological and technical content. The three IFALD donors differ in age, tissue preservation method, and disease condition (Figure 2c). In both the unintegrated PCA and default Harmony UMAP, donor C54 (a healthy adult, frozen) was clearly separated from the other two donors, suggesting that the dominant between-batch variation reflects sample-level characteristics rather than disease status per se (Figure 2b; Figure S9). Consistent with this, batch-unstable genes were enriched for gene ontology terms related to coagulation and fibrinolysis, acute-phase and inflammatory responses, and lipid and metabolic processes (Figure 2e)—processes that are likely associated with age and preservation rather than shared cell identities [46–48]. Batch-unstable genes were also enriched for mitochondrial and ribosomal genes, which are known to reflect sample quality, stress, and global transcriptional activity rather than cell identity; conversely, these genes were underrepresented among BSGs (Table S4). Notably, not all ribosomal genes were uniformly unstable: a subset with specialized regulatory functions were consistently classified as batch-stable across datasets, consistent with prior gene-wise analyses [40] (Supplementary Material; Supplementary Figure S11).

Together, these results demonstrate that BSG selection improves cell identity recovery by filtering out genes dominated by sample-specific effects or technical noise, while retaining genes that encode shared cell identity structure across batches.

### Integration scores guide hyperparameter optimization to balance under-integration and over-integration for cell identity recovery

State-of-the-art integration methods (e.g., Seurat-RPCA, Harmony, scVI, and FastMNN) produce a low-dimensional cell embedding as their integration output, and IntegrateRigor evaluates each dimension of this embedding using two complementary annotation-free metrics: BAS, which quantifies the degree of between-batch alignment, and CIS, which measures how strongly the dimension preserves biologically distinct cell identity structure. To calculate BAS and CIS, IntegrateRigor uses a Gaussian mixture model to evaluate the distribution of values in each embedding dimension. However, LIGER embeddings were not well captured by this model (see **Methods Section**), so LIGER was excluded from hyperparameter tuning and method selection analyses.

In the IFN-*β*-stimulated PBMC dataset [49] (13,999 cells, two batches), BAS and CIS were directly interpretable at the dimension level on the Harmony embedding: dimensions with high BAS showed strong batch mixing, whereas those with high CIS clearly resolved major immune cell types (Figure 2f). This confirms that the two scores capture complementary aspects of integration quality and can support exploratory analysis of individual dimensions of the integrated embedding. Building on this, IntegrateRigor aggregates the dimension-wise BAS and CIS by taking their median across all embedding dimensions, and defines the overall integration score as the sum of the two median scores. On the IFN-*β*-stimulated PBMC dataset, default Harmony showed clear under-integration, with cells remaining partially separated by batch, compromising cell identity recovery (Figure 2g). In contrast, the hyperparameter value selected by the integration score, even without BSG selection, improved batch mixing (CiLISI increased from 1.54 to 1.65) while substantially improving cell identity recovery (ARI increased from 0.73 to 0.94) (Figure 2g). Crucially, this optimization required no cell identity annotations and relied entirely on the intrinsic structure of the integrated embedding.

A key advantage of jointly scoring batch mixing and cell identity preservation is the ability to guard against over-integration, a failure mode that annotation-free batch-mixing metrics alone cannot detect. Existing annotation-free metrics such as iLISI, PCR, batch ASW, and kBET reward stronger batch mixing but do not assess whether cell identity structure is preserved. To illustrate this limitation, we analyzed a human immune cell dataset with two batches (one derived from bone marrow tissue and the other from PBMC) [17]. We then evaluated Harmony embeddings across a range of hyperparameter values. The four batch-mixing metrics showed similar trends across hyperparameters and, although kBET selected a slightly different hyperparameter value than iLISI, PCR, and batch ASW, all of these settings favored stronger batch mixing and diverged from the hyperparameter at which the IntegrateRigor integration score peaked. Specifically, along the hyperparameter path, BAS increased with the hyperparameter value, whereas CIS decreased (Figure S16), demonstrating the tradeoff between batch mixing and cell identity preservation— a consideration absent from existing batch-mixing metrics (Figure S16). At the hyperparameter selected by maximizing the integration score, cells from different batches were well aligned while major immune populations remained clearly separated (CiLISI = 1.43, cLISI = 1.16, ARI = 0.65). In contrast, the hyperparameter selected by iLISI, PCR, ASW, and kBET achieved a similar batch mixing (CiLISI = 1.40) but exhibited over-integration: several distinct cell identities became blended (cLISI increased to 1.26 and ARI dropped to 0.61) (Figure 2h). Here, a higher cLISI value indicates worse cell identity preservation.

Together, these results demonstrate that the integration score identifies hyperparameter settings that achieve a principled balance between batch mixing and cell identity preservation for reliable cell identity recovery, mitigating both under-integration and over-integration without requiring prior cell identity annotations.

### IntegrateRigor consistently improves cell identity recovery across diverse datasets and methods

To assess the generalizability of IntergrateRigor, we evaluated it across a diverse collection of integration tasks spanning multiple sources of between-batch variation: sample conditions (IFN-*β*-stimulated versus control PBMCs [49]; IFALD donors with distinct age, preservation, and disease context [44]), sequencing platforms (human immune cells [17]), developmental stages (chicken heart [14]), tissue slices (mouse brain spatial transcriptomics [50]) and species (human–mouse pancreas [51]) (summary of the datasets in Table S1). This diversity allows us to test whether IntegrateRigor can improve recovery of comparable cell identities across batches in settings that go well beyond routine single-cell batch correction, including cases where technical and biological differences make integration especially challenging.

Across all these tasks, IntegrateRigor consistently improved cell identity recovery relative to the default pipeline, as evidenced by better batch mixing (CiLISI) together with maintained or improved cell identity preservation (ARI) (Figure 3a). In the IFN-*β*-stimulated PBMC dataset, applying IntegrateRigor with scVI improved batch mixing (CiLISI increased from 1.69 to 1.78) while maintaining high cell identity recovery (ARI from 0.82 to 0.83), reflecting better mixing of stimulated and control cells without loss of cell-type structure. In IFALD, where default Harmony left cells of the same type partially separated by donor, IntegrateRigor substantially improved both batch mixing and cell identity recovery (CiLISI: 1.36 → 1.47; ARI: 0.77 → 0.88). In the chicken heart dataset, FastMNN with IntegrateRigor improved from CiLISI = 2.49 to 2.63 and ARI from 0.62 to 0.63. In the human–mouse pancreas dataset, Seurat-RPCA batch mixing increased from CiLISI = 1.88 to 2.07 while ARI remained high at 0.95. Complete metric values and UMAP visualizations for all datasets are provided in Supplementary Figures S21–S27. Together, these results demonstrate that IntegrateRigor improves cell identity recovery across a wide range of dataset scenarios, from cases where the default embedding is already reasonable to examples with pronounced under-integration.

**Figure 3:**
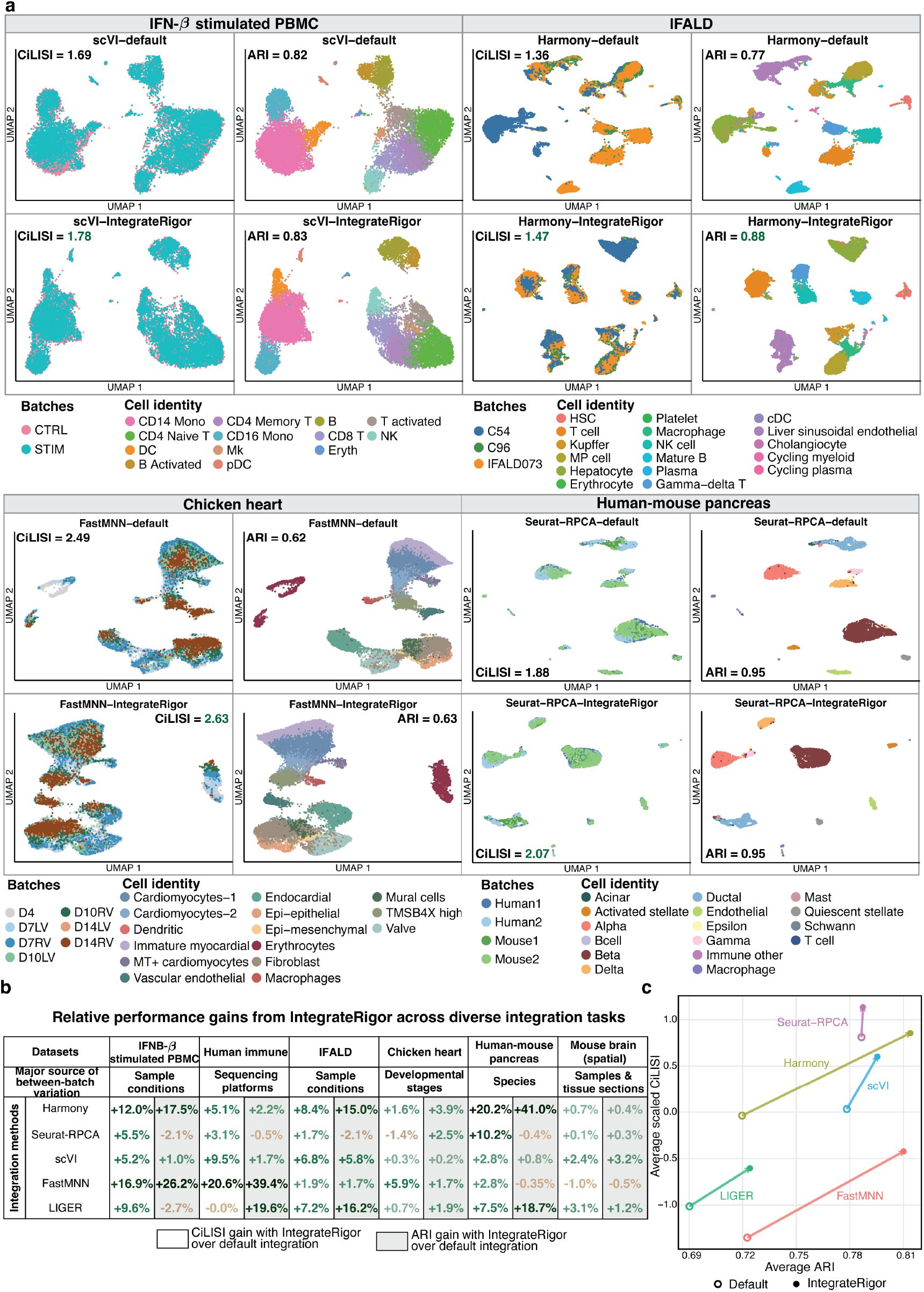
IntegrateRigor improves integration performance across diverse datasets and methods. **a**. Representative UMAPs for four datasets comparing default integration with IntegrateRigor-optimized integration. Cells are colored by batch (first column in each dataset’s subfigure) or cell identity (second column in each dataset’s subfigure). Across datasets and methods, IntegrateRigor improves batch mixing (CiLISI) while preserving or improving cell identity separation (ARI). **b**. Relative gains in CiLISI and ARI obtained by IntegrateRigor over default integration across six benchmark datasets and five integration methods. Results are grouped by the major source of between-batch variation. **c**. Method-level summary of average ARI and scaled CiLISI before and after IntegrateRigor optimization. Filled circles denote IntegrateRigor-selected solutions and open circles denote default settings.

The gains are summarized in Figure 3b, which shows the relative improvement of IntegrateRigor over default integration across methods and datasets. Batch mixing (CiLISI) improved in nearly all settings, with the largest gains in datasets with strong between-batch variation such as IFN-*β*-stimulated PBMC, IFALD, and human–mouse pancreas. The strong improvement on the human–mouse pancreas dataset shows that IntegrateRigor can also optimize cross-species integration, where the differences between batches are not limited to technical noise but include species-level transcriptomic differences. This result supports the broader applicability of IntegrateRigor: by selecting batch-stable genes and tuning integration strength, the framework can recover comparable cell identities even when datasets are separated by large biological and technical gaps. Crucially, these improvements in batch alignment were accompanied by maintained or improved ARI, confirming that increased mixing did not come at the cost of losing cell identity structure (Figure 3c). Notably, Harmony showed the largest improvement and became the best-performing method overall when optimized by IntegrateRigor, despite its moderate default performance, demonstrating how dataset-specific optimization can enable better cell identity recovery. To diagnose the sources of these improvements, we evaluated the contributions of the two stages of IntegrateRigor separately (Figure S12). BSG selection (Stage 1) accounts for most of the improvement across integration methods, whereas Harmony additionally shows a substantial gain from hyperparameter optimization (Stage 2).

We next examined whether the integration score, in addition to guiding hyperparameter selection within each method, can help distinguish among integration methods. This is inherently difficult because integration has no absolute gold standard and different metrics capture different aspects of performance [41, 52]. We therefore do not claim that IntegrateRigor selects a uniquely optimal method, but instead ask whether its ranking agrees with established annotation-dependent metrics of batch alignment and cell identity preservation.

We applied IntegrateRigor to select BSGs and optimize the hyperparameters of four integration methods, which were then ranked by the proposed integration score, excluding LIGER from this comparison. In four of the six datasets, Seurat-RPCA and Harmony were the two methods with the highest integration scores (Figure S28). One exception was the chicken heart dataset, for which Seurat-RPCA and FastMNN ranked highest, with FastMNN achieving the top integration score (Figure S28). Another exception was the IFN-*β*-stimulated PBMC dataset, for which Seurat-RPCA and scVI ranked highest (Figure S28). This pattern is broadly consistent with the overall ranking in Figure 3c, where Seurat-RPCA and Harmony achieved high CiLISI and high ARI, and with the strong cell identity separation observed for FastMNN on the chicken heart dataset (Figure S24). These observations support the integration score as a useful annotation-free proxy for evaluating method performance in cell identity recovery and batch alignment.

### IntegrateRigor uncovers cancer–immune interface niches in colorectal cancer by integrating single-cell and spatial transcriptomics data

Principled integration for reliable cell identity recovery can enable biological discoveries that remain hidden under suboptimal hyperparameter choices. We illustrate this in a colorectal cancer dataset profiled with both single-cell and spatial transcriptomics [12] (192,166 cells in total, including two MERSCOPE spatial tissue slices—PFA A11 with 86,024 cells and PFA A6 with 36,989 cells—and one 10x Genomics single-cell batch with 69,153 cells). The original study [12] used Harmony as an upstream step to obtain integrated embedding for Crescendo, a downstream count correction method, and provided cell-type annotations based on the Harmony embedding (Figure 4a). However, neither the code nor the integrated embedding used in that study was publicly released. We therefore reproduced the Harmony embedding under default settings and found it markedly different from the published embedding: the original embedding showed extensive batch mixing (Figure 4a), whereas our default Harmony embedding showed clear residual batch separation, particularly among epithelial cancer cells (Figure S29). This discrepancy highlights a common challenge: without a principled criterion, it is unclear which embedding better preserves biologically distinct cell identities.

**Figure 4:**
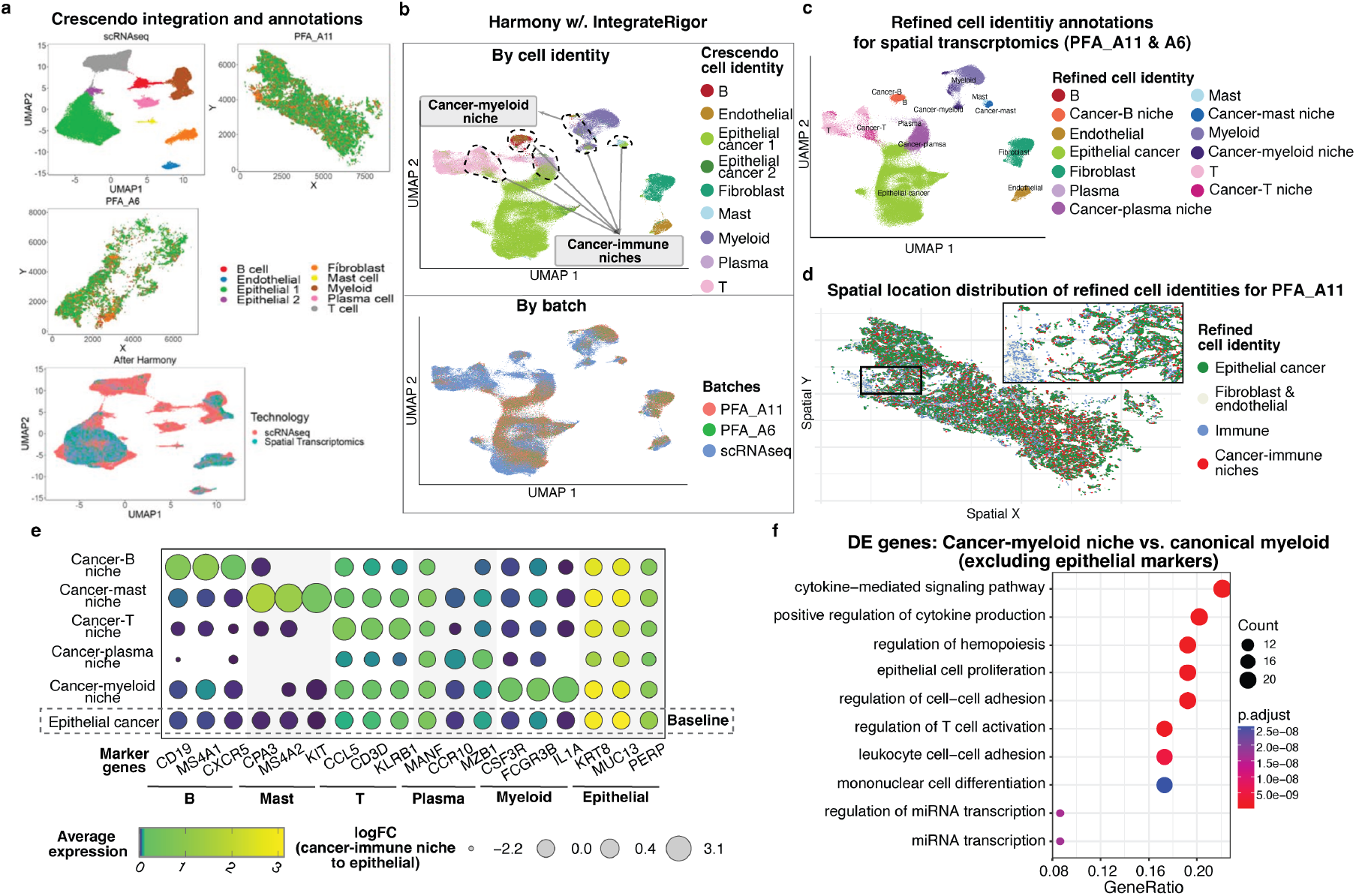
IntegrateRigor reveals unique cancer–immune interface niche in colorectal cancer data. **a**, The original Crescendo integration and annotations. The top panels show the original Crescendo embedding and spatial coordinates before integration, and the bottom panel shows the Harmony-integrated embedding colored by technology. **b**, UMAPs from Harmony integration with IntegrateRigor-selected genes, colored by batch or by Crescendo cell identity annotations. Regions where originally annotated epithelial and immune cells were intermixed are highlighted with dashed circles and were further clustered and annotated as distinct cancer–immune niches based on the IntegrateRigor embedding. **c**, Refined cell identity annotations for the spatial transcriptomics samples PFA A11 and PFA A6. Immune cells located near epithelial cancer cells were further annotated as cancer–immune niche populations, including cancer– B niche, cancer–mast niche, cancer–myeloid niche, cancer–plasma niche, and cancer–T niche. **d**, Spatial distribution of the refined cell identities in PFA A11. The main spatial map shows epithelial cancer cells, fibroblast and endothelial cells, immune cells, and cancer–immune niches. The inset highlights a zoomed-in region where cancer–immune niche cells are spatially intermingled with epithelial cancer cells. **e**, Dot plot of canonical marker genes across refined cell identity groups. Dot color represents average expression, and dot size represents the log fold change of each cancer– immune niche population relative to epithelial cancer cells. Cancer–immune niche groups retained expression of their corresponding immune markers while showing epithelial marker expression at levels comparable to epithelial cancer cells. **f**, Gene Ontology enrichment analysis of differentially expressed genes comparing the cancer-myeloid niche with myeloid cells after excluding epithelial marker genes. Enriched pathways include cytokine-mediated signaling, regulation of cytokine production, regulation of cell–cell adhesion, and T cell activation, supporting the biological distinction of the cancer-myeloid niche from conventional myeloid cells.

Applying IntegrateRigor to this dataset yielded an optimized Harmony embedding that differed from both the original and the default result, achieving a better balance between batch mixing and cell identity preservation (Figure 4a). When we examined whether the cell-type annotations from the original study remained consistent with the optimized embedding, we found substantial discordance: a subset of cells originally annotated as epithelial cancer cells clustered among immune populations in the IntegrateRigor embedding. This motivated us to perform independent cell-type annotation on the optimized embedding using Louvain clustering and marker gene expression.

In our reannotation, cells that shifted from epithelial cancer cells to immune cells were mapped back to their physical locations in the spatial transcriptomics data (Figure 4b). These cells localized specifically to the boundary between canonical epithelial cancer cells (annotated as epithelial cancer in both the original and revised annotations) and canonical immune cells (annotated as immune in both annotations). We refer to these spatially localized boundary populations as *cancer–immune interface niches*.

We confirmed their identity as interface populations by differential expression analysis using observed gene expression from a single batch before integration, to avoid integration artifacts (Figure 4c). When compared with canonical epithelial cancer cells, each niche was enriched for markers of its corresponding immune cell type; when compared with canonical immune cells, each niche retained strong epithelial marker expression (*KRT8, MUC13, PERP*). This dual signature— immune-like relative to epithelium, epithelial-like relative to immune—defines the interface niche identity. Specifically, the cancer–B niche expressed *CD19, MS4A1*, and *CXCR5*; the cancer– mast niche expressed *CPA3, MS4A2*, and *KIT* ; the cancer–T niche expressed *CCL5, CD3D*, and *KLRB1*; the cancer–plasma niche expressed *MANF, CCR10*, and *MZB1*; and the cancer–myeloid niche expressed *CSF3R, FCGR3B*, and *IL1A*. Such mixed transcriptional profiles are expected in spatial transcriptomics data, where direct cell–cell contact, imperfect cell segmentation, and local transcript contamination can all contribute.

Functional enrichment analysis of genes differentially expressed between each niche and its corresponding canonical immune population (excluding shared epithelial markers) revealed immune cell-type-specific programs consistent with active roles at the cancer–immune interface (Figure 4d; Figure S30) [53–55]. The cancer–myeloid niche was enriched for “cytokine-mediated signaling pathway,” “positive regulation of cytokine production,” “leukocyte cell–cell adhesion,” and “regulation of T cell activation,” consistent with a program of immune communication and local coordination in the tumor microenvironment [56, 57]. The cancer–B niche was dominated by antigen-processing and antigen-presentation terms, consistent with a state specialized for antigen handling at the epithelial cancer interface [58–60]. The cancer–T niche was enriched for leukocyte-mediated cytotoxicity, interleukin-10 production, and cell–cell adhesion, suggesting an immunoregulatory T-cell state shaped by epithelial contact [57, 61]. The cancer–plasma niche was enriched for antigen-presentation and activated immune-response programs beyond canonical plasma-cell function [62]. The cancer–mast niche was enriched for immunological memory formation and antigen-presentation-related terms, suggesting involvement in broader adaptive immune regulation [63–65]. Across all niches, the enriched programs reflect the characteristic functions of the corresponding immune cell type, consistent with their positions at the cancer– immune interface.

These interface niches were not detectable under either the original or the default Harmony embedding. The original embedding merged them into a broad cluster, masking their intermediate identity and leading to their annotation as epithelial cancer cells only. The default Harmony embedding failed to align them consistently across batches, preventing their recognition as specific types of niches. In contrast, the IntegrateRigor-optimized embedding revealed these cancer–immune interface niches and supported their interpretation through spatial localization, dual cancer–immune marker signatures, and immune cell-type-specific functional enrichment. This example demonstrates that the choice of integration hyperparameters is not merely a technical detail: by optimizing integration for cell identity recovery, IntegrateRigor can reveal cell populations of biological interest that would otherwise remain obscured.

## Discussion

IntegrateRigor reframes data integration from a heuristic preprocessing step into a dataset-driven, statistically guided optimization problem for cell identity recovery. Rather than introducing a new integration method, it provides a method-agnostic framework for improving existing methods through annotation-free batch-stable gene selection and hyperparameter tuning. Across diverse single-cell and spatial transcriptomics tasks, this strategy improved cell identity recovery by better balancing batch mixing and cell identity preservation, reducing both under-integration and over-integration, and revealing biologically meaningful structure that was not apparent under default settings.

Our results highlight two underappreciated points about integration practices. First, gene selection is itself a major determinant of cell identity recovery—a point that receives far less attention than the choice of integration method, even though the selected genes define the information the method operates on when aligning cells across batches. By identifying BSGs and excluding genes dominated by between-batch variation, IntegrateRigor improves the ability of diverse methods to recover comparable cell identities across batches. Second, integration performance depends not only on the method chosen but also on how that method is configured for a given dataset. Harmony showed only moderate performance with default settings but became one of the best-performing methods after IntegrateRigor-based optimization. This illustrates that method rankings can shift substantially when gene selection and hyperparameter tuning are made more principled. More broadly, these findings argue for moving beyond static, one-size-fits-all integration pipelines and toward dataset-specific optimization, where the central question is how the method configuration best supports cell identity recovery for a particular dataset.

The quality of the integrated embedding has consequences that extend well beyond the integration step itself. Because cell identity recovery is a prerequisite for any cell-identity-based downstream analysis, the reliability of the integrated embedding determines the quality of all analyses that follow. For example, Crescendo [12] and CellANOVA [30] both take the integrated embedding directly as input to perform count-level batch correction and cell-type-specific differential expression analysis under particular experimental designs, respectively. Because their cell-state representations and decompositions are estimated from the integrated embedding, the reliability of their outputs depends directly on the quality of cell identity recovery in that embedding. An under-integrated embedding retains residual batch structure that can confound cell-state estimation; an over-integrated embedding blurs biologically distinct states, causing cell-type mixing that corrupts cell-type-specific analyses. Providing these downstream methods with a well-calibrated embedding—one that supports reliable cell identity recovery across batches—is therefore a prerequisite for accurate downstream inference.

It is worth clarifying the relationship between IntegrateRigor and CellANOVA [30]. CellANOVA recovers biological signals associated with within-cell-identity between-batch variation that is orthogonal to the experimental design—variation that may be inadvertently removed during integration. In that sense it addresses a form of over-integration, but its aim is distinct from ours. IntegrateRigor addresses the upstream task of cell identity recovery: reliably aligning cells with the same identity across batches before any downstream analysis is performed. For this cell identity recovery task, within-cell-identity between-batch variation is treated as residual under-integration to be corrected, not as signal to be recovered. Moreover, CellANOVA still depends on an existing integrated embedding to estimate its cell-state encoding, so its performance remains sensitive to the quality of that upstream embedding. IntegrateRigor addresses this dependency by providing an optimized embedding as input.

The importance of integrated embedding for cell identity recovery was directly illustrated in the colorectal cancer application. Applying IntegrateRigor to obtain an optimized Harmony embedding enabled more reliable cell identity recovery, which in turn revealed previously unidentified cancer– immune interface niches in which distinct immune cell populations showed transcriptional and spatial signatures consistent with direct interaction with malignant epithelial cells. Without careful gene selection and parameter tuning, the default Harmony embedding left residual batch structure that prevented coherent cell identity recovery, whereas the embedding from the original study overcorrected and obscured these interface populations. This example illustrates how integration optimization for cell identity recovery directly shapes which cell populations can be distinguished.

Several extensions could further strengthen this framework. First, the current formulation focuses on cell identity preservation as the primary objective of integration optimization, treating between-batch variation as the target of correction. A natural next step is to incorporate known sample-level covariates (e.g., treatment condition, disease status, or developmental stage) to distinguish generic batch effects from structured biological variation of interest. With this experimental design information, the same optimization principles could be extended beyond the cell identity recovery task to directly optimize the integrated embedding for downstream analyses such as those performed by Crescendo and CellANOVA: for example, optimizing integration that maximizes power for condition-specific differential expression while controlling for batch. This extension reflects the natural order of analysis: first obtain reliable cell identities, then leverage them for cell-identity-based inference. Second, the current IntegrateRigor does not incorporate spatial information when evaluating the integration of spatial transcriptomics data. Extending IntegrateRigor to settings where all batches are spatial transcriptomics would therefore require an additional metric beyond batch mixing (BAS) and cell identity preservation (CIS). For spatial transcriptomics, a well-integrated embedding should not only align cells with the same identity across batches, but also preserve spatial organization within each tissue section: spatially neighboring cells or spots are expected to be nearby in the integrated embedding. Third, IntegrateRigor relies on two modeling assumptions: a negative binomial mixture model (NBMM) for per-gene expression counts and a Gaussian mixture model (GMM) for per-dimension values in the integrated embedding; future work could explore more flexible latent models or nonparametric density estimators that better capture the geometry of embeddings that deviate from these assumptions, such as those produced by LIGER. Fourth, we have so far assumed that a single reference batch contains all relevant cell identities, an assumption that is increasingly reasonable given the availability of large atlases that can serve as references. In settings where each batch contains only a subset of cell identities, a natural extension would be to construct a synthetic reference by merging and aligning mixture-model components across batches, rather than relying on a single reference batch; this will require methodological development in future work.

More broadly, our findings suggest that future benchmarks should evaluate integration methods not only under default settings but also after dataset-specific optimization, as default performance may substantially underestimate the quality achievable for cell identity recovery with principled configuration. The same principles underlying IntegrateRigor may also extend to cross-modality integration, where preserving shared cell identity across different feature spaces presents additional challenges, and to semi-supervised settings in which partial reference annotations are available, analogous to reference-guided approaches such as scANVI [28, 66].

## Methods

We first consider a two-batch setting consisting of a reference batch and a query batch, and subsequently extend the framework to multiple batches. The reference batch serves as an anchor for shared cell identity structure and is assumed to contain a more diverse set of cell identities, while the query batch may contain only a subset of cell identities.

Let the gene expression count matrix of the reference batch be denoted by 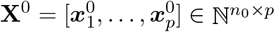 and that of the query batch by 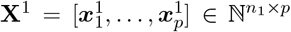, where *n*_0_ and *n*_1_ denote the numbers of cells in the reference and query batches, respectively, and *p* denotes the number of genes.

### Batch-stable gene (BSG) selection for data integration

#### Statistical modeling for gene expression in reference and query batches

We focus on a single gene and suppress the gene index for notational simplicity. Let 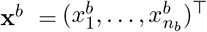 denote the expression counts in batch *b* ∈ {0, 1}, with corresponding library sizes 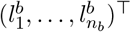. Here, *b* = 0 refers to the reference batch, and *b* = 1 refers to the query batch.

To identify genes that preserve cell identity across batches, we model gene expression using a negative binomial (NB) mixture model, with mixture components representing distinct cell identities. This model is motivated by extensive prior evidence that the NB distribution effectively models overdispersed gene-expression count data within a homogeneous cell identity [67]. Validation of this model assumption is provided in **Methods Subsection “Validation of the statistical models underlying IntegrateRigor”**. Specifically, we assume that for any given gene

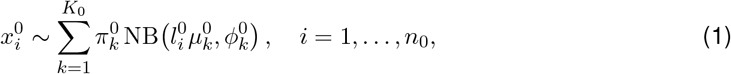

and

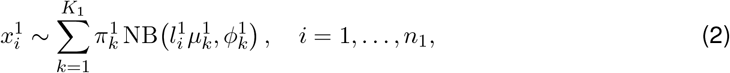

where *K*_1_ ≤ *K*_0_. Here, 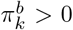 denotes the mixing proportion of component *k* in batch *b* ∈ {0, 1}, with 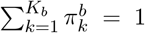, and 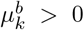 and 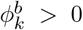 denote the corresponding mean and dispersion parameters for that NB component, respectively. We assume that these component-specific parameters are distinct across mixture components; that is,

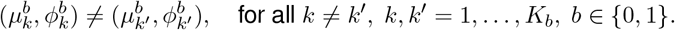

Intuitively, if a gene reflects cell identity rather than between-batch variation, its expression pattern across cell identities should be preserved across batches, even when the proportions of those cell identities differ. In this case, the gene expression distribution in the query batch is assumed to share a subset of the mixture components in the reference batch, while allowing only the mixing proportions to differ.

Specifically, in the absence of between-batch variation, the mixture components 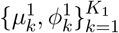 in the query batch model (2) is a subset of the mixture components 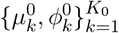 in the reference batch model (1). Then the gene expression distribution of the query batch can be re-parameterized using 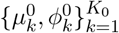, while allowing only the component proportions to vary:

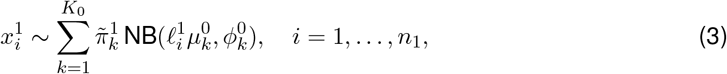

which we refer to as the *reference-constrained model*. Here, 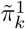 denotes the mixing proportion of the reference component *k* in the reference-constrained model for the query batch. We distinguish it from 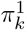, which refers to the mixing proportion of the query component *k* in model (2), and the reference components and query components may not share the same indices.

In contrast, between-batch variation induces shifts in the component parameters, such that 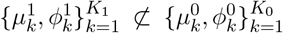. Consequently, the query distribution can no longer be adequately explained by the reference mixture structure alone and instead requires the *full model* (2). We therefore define the BSS for the gene based on how well the query batch can be explained by the reference-constrained model relative to the full model.

#### Batch stability score (BSS) definition

To distinguish these two scenarios, we compare two models, the reference-constrained model (3) and the full model (2), for the query batch, both fitted using the Expectation–Maximization (EM) algorithm [68]. Under the reference-constrained model (3), the component parameters 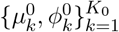 are first estimated from the reference batch via EM and then treated as fixed when evaluating the query batch, with only the mixing proportions 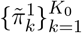 re-estimated by EM. Under the full model (2), all parameters 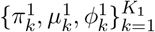 are estimated directly from the query batch using EM.

Let 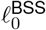 denote the log-likelihood of the reference-constrained model and 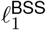 of the log-likelihood of the full model, both evaluated based on the parameter estimates from the query batch. Then we define the batch stability score (BSS) as

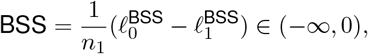

which takes a negative value as 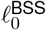 is strictly smaller than 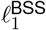 as the parameter space of the reference-constrained model is a strict subspace of that of the full model.

A high (less negative) BSS indicates that the reference-constrained model explains the query data well, suggesting that the gene’s expression pattern transfers between batches and is therefore batch-stable. Conversely, a low (more negative) BSS indicates that the full model provides a substantially better fit, suggesting between-batch variation. BSGs are selected based on the empirical distribution of BSS values using a data-adaptive threshold determined by an elbow criterion [45].

### Integration score for a given dataset and integration method

#### Statistical modeling for integrated embedding

Let 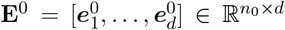 and 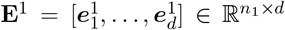 denote the low-dimensional embeddings of the reference and query batches from an integrated embedding produced by a given integration method, where 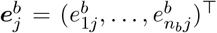 denotes the *j*th embedding dimension for batch *b* ∈ {0, 1}, where *b* = 0 denotes the reference batch, and *b* = 1 denotes the query batch.

For each embedding dimension *j* = 1, …, *d*, we assume that

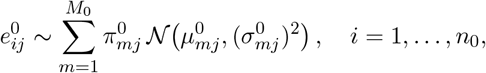

and

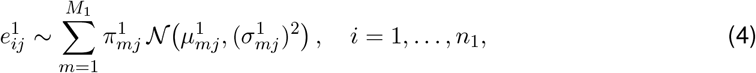

where *M*_1_ ≤ *M*_0_. Here, 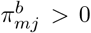 denotes the mixing proportion of component *m* in embedding dimension *j* for batch *b* ∈ {0, 1}, with 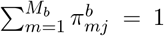, and 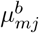 and 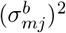 denote the corresponding mean and variance parameters for that Gaussian component, respectively. Given our observation that the Gaussian mixture model reflects distinct cell identities in some dimensions of the integrated embedding (see **Methods Subsection “Validation of the statistical models underlying IntegrateRigor.”**), we assume that the mixture components in this model correspond to cell identities and that the component-specific parameters are distinct; that is,

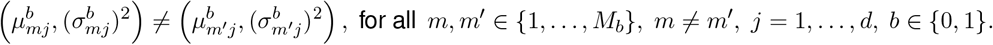

#### Batch alignment score (BAS) definition

The BAS quantifies batch mixing in the integrated embedding space. For each embedding dimension *j* = 1, …, *d*, following the same rationale as the BSS definition, we compare a reference-constrained model,

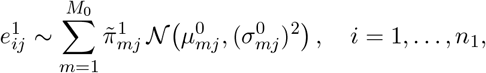

with the full model (4) for the query batch embeddings in the same dimension. Under the reference-constrained model, the component parameters 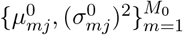 are estimated from the reference batch and then treated as fixed when evaluating the query batch, with only the mixing proportions 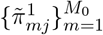 re-estimated. Under the full model, all parameters 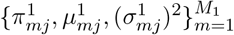 are estimated directly from the query batch.

Let 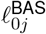 denote the log-likelihood of the reference-constrained model and 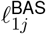denote the log-likelihood of the full model, both evaluated on the query embeddings in dimension *j*. We define the dimension-wise BAS as

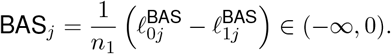

The overall BAS is defined as the median across all embedding dimensions:

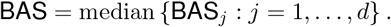

A higher (less negative) BAS indicates that the reference-constrained model explains the query embeddings well, suggesting better alignment between batches and more effective removal of between-batch variation. Conversely, a lower (more negative) BAS indicates that the full model provides a substantially better fit, suggesting residual between-batch variation and potential under-integration.

#### Cell identity score (CIS) definition

The CIS quantifies the extent to which biologically distinct cell identities are preserved in the integrated embedding. Intuitively, an embedding dimension that captures cell identities should exhibit a multimodal distribution, whose modes (i.e., mixture components) correspond to distinct cell identities [10]. In contrast, if an embedding dimension exhibits a unimodal distribution, it does not reveal cell identities.

Thus, for each embedding dimension *j*, we compare a Gaussian mixture model with a single Gaussian model, both fitted within each batch. Let 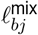 denote the log-likelihood of the Gaussian mixture model and 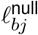 denote the log-likelihood of the single Gaussian model evaluated on batch *b* = 0, 1 and dimension *j*. We define the dimension-wise CIS as

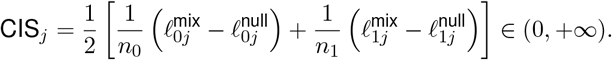

The overall CIS is defined as the median across all embedding dimensions:

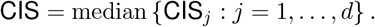

A higher CIS indicates that the Gaussian mixture model explains the embedding values well across dimensions, suggesting stronger preservation of cell identity structure, whereas a low CIS suggests loss of cell identities and potential over-integration.

#### Integration score definition

The dataset-level integration score is defined as

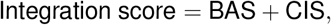

where BAS penalizes under-integration, and CIS penalizes over-integration. Hence, the integration score balances the trade-off between batch mixing and cell identity preservation, providing a principled, annotation-free criterion. Maximizing the integration score, IntegrateRigor optimizes the hyperparameter setting of a given integration method on a given dataset.

#### Hyperparameter for integration methods

On the six integration datasets we analyzed in the **Subsection “IntegrateRigor consistently improves integration across diverse datasets and methods**,**”** we evaluated the following hyperparameters and values for each integration method.

For Harmony, we performed a grid search over two hyperparameters: `theta` ∈ {2, 4, 8}, which controls the strength of the diversity clustering penalty and `nclust` ∈ {80, 100, 120}, which controls the number of clusters used in soft clustering. This grid search resulted in 3 × 3 = 9 parameter settings for each dataset. For Seurat-RPCA, we searched across the hyperparameter, `k.weight` ∈ {20, 60, 80, 100, 140}, which determines the number of neighbors considered when weighting anchors and is associated with the degree of between-batch variation removal. For scVI, we searched across the hyperparameter, `n_hidden` ∈ {64, 128, 256}, which specifies the number of nodes in each hidden layer of the neural network. For FastMNN, we searched across the hyperparameter, `k` ∈ {10, 20, 60, 100}, which specifies the number of nearest neighbors used for mutual nearest-neighbor matching across batches.

### Generalization to multiple batches

Suppose there are *B* > 1 query batches in addition to one reference batch. Let

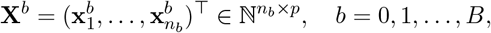

denote the gene-count matrix for batch *b*. Here, *b* = 0 refers to the reference batch, whereas *b* = 1, …, *B* refer to the query batches. For each query batch *b* ∈ {1, …, *B*}, comparing it with the reference batch yields a gene-level BSS^*b*^. We then define the overall BSS for each gene by averaging across query batches:

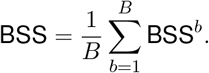

Next, let

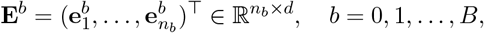

denote the corresponding low-dimensional embedding matrix. For each query batch *b* ∈ {1, …, *B*}, comparing it with the reference batch yields a dimension-level 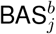 for embedding dimension *j*. We then define the overall BAS_*j*_ for each dimension by averaging across query batches:

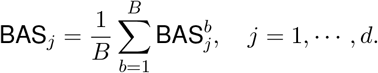

The per-dimension CIS generalizes naturally to the multi-batch setting by averaging the per-batch log-likelihood improvement over all batches, including the reference batch:

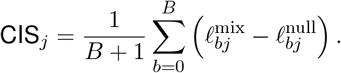

Finally, the dataset-level integration score is defined as the sum of the median BAS and median CIS across embedding dimensions:

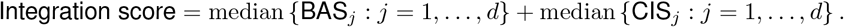

### Validation of the statistical models underlying IntegrateRigor

To validate the modeling assumptions underlying IntegrateRigor, we examined the goodness-of-fit of the two probabilistic models: the NB mixture model (NBMM) for gene-level expression counts and the Gaussian mixture model (GMM) for each dimension of the integrated embedding. From the intestinal failure-associated liver disease (IFALD) dataset [44], we selected three donors comprising 32,397 genes and 15,895 cells. For embedding-level validation, we applied five integration methods (Seurat-RPCA, Harmony, scVI, FastMNN, and LIGER) with default settings to integrate the three batches and evaluated the GMM fit to the resulting integrated embedding (dimension-wise) for each method. For gene-level validation, we applied the NBMM to one batch before integration, as an illustrative example.

We quantified the goodness-of-fit using the Kolmogorov–Smirnov (KS) statistic [69], defined as the maximum absolute difference between the empirical distribution function and the fitted cumulative distribution function, to measure the magnitude of distributional discrepancy. We did not use the p-value to qualitatively assess goodness-of-fit, because it is well known that for large sample sizes—such as those in large-scale single-cell datasets—the KS p-value tends to become vanishingly small, so that even minor and practically negligible deviations from the fitted distribution can lead to rejection of the model [70].

For modeling the per-gene expression counts, the observed-versus-fitted histograms show that the NBMM closely matches the count distributions for representative genes in IFALD. This is consistent with extensive prior evidence that the NB distribution provides an effective model for overdispersed gene-expression count data in homogeneous cell populations (cells with the same underlying identity) [67]. Across the displayed examples, most genes have very small KS statistic values, often near zero and generally below 0.02 (Figure S2). This pattern extends beyond the examples shown. Across 2000 HVGs, the distribution of KS statistic values is concentrated near zero, with only a modest upper tail and a limited number of outliers (Figure S1a). Overall, these observations support the NBMM as a reasonable model for gene-level distribution modeling.

For modeling the per-dimension integration embedding values, Gaussian modeling is motivated both empirically and theoretically: classical results on linear dimension reduction imply that many low-dimensional embeddings of high-dimensional data are approximately Gaussian [71]. To evaluate this GMM assumption empirically, we fitted a GMM to each of the first five dimensions of the integrated embedding produced by each of the five integration methods and assessed the goodness-of-fit using the KS statistic (Figure S3). For scVI, Harmony, Seurat-RPCA, and FastMNN, the KS statistic values are consistently small across the first five dimensions, generally on the order of 0.004–0.016, indicating that their dimension-wise embedding values are well approximated by the GMM. The summary boxplot across dimensions shows the same trend, with the KS statistic values concentrated near zero (Figure S1b).

In contrast, the integrated embedding of LIGER exhibits substantially larger KS values, around 0.223–0.332 in the first five dimensions, indicating a poorer goodness-of-fit for GMM relative to the integrated embeddings of the other four methods. This result is consistent with the fact that LIGER is based on non-negative matrix factorization, which tends to produce embeddings with nonnegative, skewed, and less Gaussian-like distributions. As a result, we excluded LIGER from hyperparameter tuning and method selection analyses that rely on our integration score, which assumes a GMM.

We also validated the NBMM and GMM modeling assumptions on a spatial transcriptomics dataset. In the MERFISH mouse brain dataset [50], we observed patterns consistent with those in the scRNA-seq datasets: the NBMM achieved small KS statistics across genes, with a median KS statistic below 0.01, and the GMM fit the integrated embeddings well for most integration methods, with near-zero KS statistics across embedding dimensions. The only exception was again LIGER, for which the median KS statistics across embedding dimensions exceeded 0.2 (Figures S4–S6).

Taken together, these results support the two core modeling assumptions in IntegrateRigor: the NBMM provides a good fit to per-gene expression counts, and the GMM provides a reasonable approximation for per-dimension values in the integrated embedding. In practice, these distributional assumptions should be checked before applying IntegrateRigor, to ensure that the NBMM and GMM remain appropriate and that IntegrateRigor results are valid for a given dataset.

### Stability of IntegrateRigor

The NBMM was fitted using a custom EM algorithm, whereas the GMM was fitted using the `Mclust` function in the R package `mclust`. The key parameters for IntegrateRigor are the number of mixture components, i.e., *K*_0_, *K*_1_, *M*_0_, and *M*_1_. However, for the mixture-model fitting, as long as the specified number of mixture components is no smaller than the true number of underlying mixture components, our scores (BSS, BAS, and CIS) are expected to remain stable across different choices of *K*_0_, *K*_1_, *M*_0_, and *M*_1_ [72]. By default, the number of mixture components was set to 5 for the NBMM and GMM (*K*_0_ = *K*_1_ = *M*_0_ = *M*_1_ = 5). Although the total number of cell identities (e.g., cell types) in a dataset is often larger than 5, the distribution for a single gene or a single embedding dimension typically does not exhibit many distinct modes, because each cell identity is defined jointly by multiple genes and multiple embedding dimensions. Empirically, we observed that the number of modes per gene or embedding dimension is modest, and setting the number of mixture components to 5 is sufficient in practice.

We further evaluated the stability of IntegrateRigor with respect to the specified number of mixture components. Specifically, we examined the stability of the BSS, the selected BSGs, and the integration score across varying numbers of mixture components: *K*_0_ = *K*_1_ = *M*_0_ = *M*_1_ ∈ {3, 4, 5, …, 10}. Regarding the gene-wise NBMM fitting using our custom EM algorithm implementation, we vary the number of EM iterations from 10, 20, to 50, with 10 iterations used as the default. Across these settings, the resulting BSS values, whose calculation relies on NBMM fitting, were consistently highly correlated with those obtained under the default setting (with the Spearman correlation close to 1), and the selected BSGs showed substantial overlap with the default BSG set (with the Jaccard index for set similarity close to 1). Regarding the embedding-level GMM fitting under varying numbers of mixture components, the integration score consistently selected the same optimal hyperparameter. These results suggest that IntegrateRigor is a stable procedure. Further details are in Supplementary Figures S17–S20.

We also evaluated the stability of IntegrateRigor to reference-batch selection. IntegrateRigor requires a reference batch that contains all cell identities present in the dataset; in some datasets, multiple batches may satisfy this requirement and can therefore serve as candidate reference batches. To assess stability to this choice, we used the IFALD dataset, in which all three batches (C54, C96, and IFALD073) contain nearly all major cell identities and can therefore each serve as the reference batch; the only exception is C54, which lacks cells annotated as cDCs. Cell-type compositions within each batch are summarized in Table S2. By default, IntegrateRigor automatically selected IFALD073 as the reference batch, as it has the largest number of Louvain clusters across resolutions. We then repeated BSG selection and integration score calculation using each of the remaining batches as the reference. The selected BSGs were highly consistent across reference-batch choices, with pairwise Jaccard similarities greater than 0.98 (Table S3). Integration scores showed broadly consistent patterns across hyperparameter values regardless of reference-batch choice, with the most similar patterns observed for C96 and IFALD073 as references (Spearman correlation 0.945); moreover, all reference-batch choices led to the same hyperparameter values being selected (Figure S8). Together, these results indicate that IntegrateRigor is robust to the choice of reference batch when multiple batches are eligible.

### Annotation-based evaluation metrics

#### Cell-type-aware integration local inverse Simpson’s index (CiLISI)

The integration local inverse Simpson’s index (iLISI) is a local neighborhood-based metric for batch mixing. For each cell, iLISI measures the diversity of cells from different batches among its nearest neighbors in the integrated embedding. A higher iLISI value indicates that the cell’s local neighborhood contains a more balanced mixture of batches, whereas a lower value indicates that the neighborhood is dominated by a subset of batches.

To evaluate batch mixing while accounting for cell identity preservation, we used a cell-type-aware version of iLISI [17, 27, 41], termed as CiLISI. For each cell type, we restricted the integrated embedding to cells annotated as that cell type and computed iLISI using batch labels within this cell subset. Thus, each cell’s iLISI value reflects batch diversity among its nearest neighbors from the same cell type, avoiding confounding from differences of distinct cell types.

We then averaged the cell-level values within each cell type to obtain a cell-type-specific iLISI. These cell-type-specific iLISIs were further averaged across cell types, weighted by cell-type proportions, to obtain the final CiLISI score. Higher CiLISI indicates better mixing of batches within the same cell type.

#### Adjusted Rand index (ARI)

To evaluate preservation of cell identity structure, we used the ARI between unsupervised clusters obtained from the integrated embedding and known cell identity labels [17, 41]. For each integrated embedding, we performed Seurat Louvain clustering over a range of resolution parameters from 0.1 to 1.0, in increments of 0.05. At each resolution, we computed the ARI between the resulting cluster assignments and the known cell identity labels. The maximum ARI across all tested resolutions was taken as the final ARI score for that embedding. This procedure avoids dependence on a single clustering resolution and captures the extent to which the integrated embedding reflects the cell identity structure.

#### Cell-type local inverse Simpson’s index (cLISI)

For each cell, cLISI measures the diversity of cell-type labels among its nearest neighbors. A lower cLISI value indicates that the local neighborhood is dominated by cells of the same cell type, reflecting better preservation of cell identity structure. A higher cLISI value indicates stronger local mixing of different cell types, which may suggest loss of cell type separation or over-integration.

We first computed LISI using cell-type labels across all cells, without conditioning on batch, and refer to these scores as cell-level cLISI values. Each cell’s cLISI value reflects the local diversity of cell-type labels among its neighboring cells, allowing cLISI to detect the mixing of distinct cell types across batches and thereby capture potential over-integration. We then averaged the cell-level cLISI values across cells to obtain the dataset-level cLISI value.

## Data availability

The IFNB dataset can be accessed through the `SeuratData` package using `SeuratData::LoadData (“ifnb”)`. The human immune dataset can be accessed at https://doi.org/10.6084/m9.figshare.12420968. The IFALD dataset can be accessed at https://cellxgene.cziscience.com/collections/ff69f0ee-fef6-4895-9f48-6c64a68c8289. The chicken heart dataset can be accessed at GSE149457. The colorectal cancer spatial transcriptomics and single-cell RNA-sequencing dataset can be accessed at https://zenodo.org/records/14602110. The mouse brain spatial transcriptomics dataset was obtained from the Allen Brain Cell Atlas using the `abc_atlas_access` package. Specifically, we used three Zhuang-ABCA slices, `Zhuang-ABCA-1.079`, `Zhuang-ABCA-2.037`, and `Zhuang-ABCA-3.010`, from the `Zhuang-ABCA-1`, `Zhuang-ABCA-2`, and `Zhuang-ABCA-3` datasets, respectively. The data can be accessed through the Allen Brain Cell Atlas data-access portal and Python package at https://alleninstitute.github.io/abc_atlas_access/.

## Code availability

The IntegrateRigor R package is publicly available at https://github.com/zhiqianZ/IntegrateRigor. And the analysis scripts are available at https://github.com/zhiqianZ/IntegrateRigor_analysis.

## Acknowledgments

The authors appreciate the comments and feedback from the Junction of Statistics and Biology members (https://jsb-lab.org).

## Author contributions

Z.Z., C.W., and J.J.L. conceived and designed the method. Z.Z., C.J., and C.W. performed the data analysis. Z.Z., C.W., Z.R., and J.J.L. interpreted the results and prepared the figures. Z.Z., C.W., C.J., Z.R., and J.J.L. drafted and revised the manuscript. J.J.L. and C.W. supervised the research and manuscript preparation.

