## Supplementary for "IntegrateRigor: annotation-free integration optimization for cell identity recovery reveals cancer–immune interface niches"

#### Contents

**1 Dataset Summary** **3**

**2 Empirical Validation of IntegrateRigor** **3**

2.1 Empirical Validation on scRNAseq data . . . . . . . . . . . . . . . . . . . . . . . . 3

2.2 Empirical Validation on MERFISH Data . . . . . . . . . . . . . . . . . . . . . . . . 5

**3 Pearson correlation between different integrated embedding dimensions** **7**

**4 Reference-batch choice stability** **8**

**5 Genes contributing to batch effect in the unintegrated PCA overlaps with the batch-unstable genes** **9**

**6 Batch stability score correlates with annotation-dependent GTE scores** **10**

**7 IntegrateRigor identifies ribosomal genes more frequently as batch-unstable genes** **11**

**8 Metrics evaluation for different stages of IntegrateRigor** **13**

**9 Figures with complete legends for main figures** **13**

**10 Harmony parameters selected by existing metrics** **15**

**11 IntegratRigor sensitivity and convergence to the IntegrateRigor hyperparameters** **15**

**12 Complete IntegrateRigor results** **18**

### 1 Dataset Summary

**Table S1:** Summary of datasets used in this study.

| Dataset | # Cells | # Batches | # Cell identities | # Genes | Reference |
|---|---|---|---|---|---|
| IFN-$\beta$ stimulated PBMC | 13999 | 2 | 13 | 14053 | [1] |
| IFALD | 15895 | 3 | 17 | 32397 | [2] |
| Immune | 32484 | 9 | 16 | 12303 | [3] |
| Chicken heart | 22315 | 7 | 15 | 24356 | [4] |
| Human-mouse pancreas | 5547 | 4 | 17 | 11820 | [5] |
| Mouse brain spatial | 137442 | 3 | 34 | 1122 | [6] |
| CRC (scRNA-seq & spatial) | 192166 | 3 | 9 | 477 | [7] |

### 2 Empirical Validation of IntegrateRigor

#### 2.1 Empirical Validation on scRNAseq data

To empirically assess whether the distributional assumptions used in IntegrateRigor are reasonable for real single-cell data, we evaluated the goodness-of-fit of the two key modeling components on the IFALD dataset. First, we examined the negative binomial mixture model (NBMM) used to model gene expression counts. For the 2,000 highly variable genes, we compared the observed nonzero count distributions with the fitted distributions and quantified the discrepancy using the Kolmogorov–Smirnov (KS) statistic.

Second, we evaluated the Gaussian mixture model (GMM) assumption used for integrated embedding dimensions. For each integration method, we fitted a GMM with $K = 5$ components to each embedding dimension and compared the fitted density with the empirical distribution. Across Seurat-RPCA, FastMNN, Harmony, and scVI, the fitted GMMs closely matched the observed embedding distributions, with generally small KS statistics across dimensions. In contrast, LIGER showed substantially larger KS statistics, indicating poorer agreement with the GMM assumption for its embedding space. These results suggest that the modeling assumptions underlying IntegrateRigor are broadly compatible with commonly used integration outputs, while also allowing the framework to detect cases where an embedding distribution deviates from the assumed mixture structure.

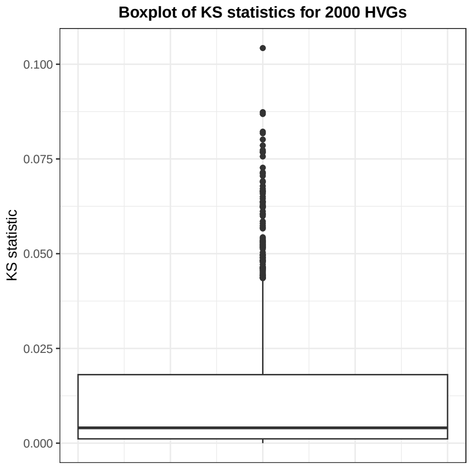

**(a)** NBMM KS statistics

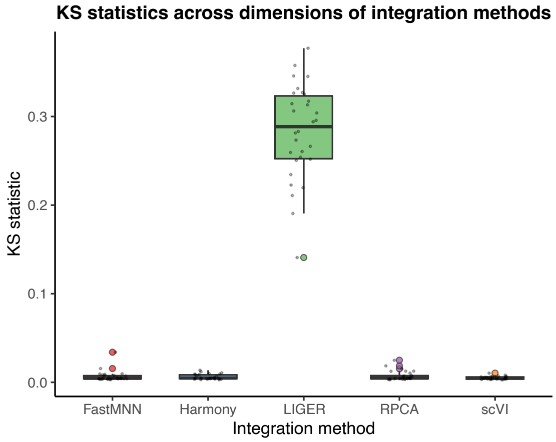

**(b)** GMM KS statistics

**Figure S1: Goodness-of-fit assessment using KS statistics on the IFALD dataset.** Left panel: Distribution of KS statistics for 2,000 highly variable genes fitted by the negative binomial mixture model (NBMM), showing generally small distributional discrepancies between the observed and fitted gene expression distributions. Right panel: Distribution of KS statistics across embedding dimensions for Gaussian mixture model (GMM) fits to integrated embeddings from different integration methods. Most methods show small KS statistics, whereas LIGER exhibits substantially larger KS values, indicating poorer compatibility with the GMM assumption.

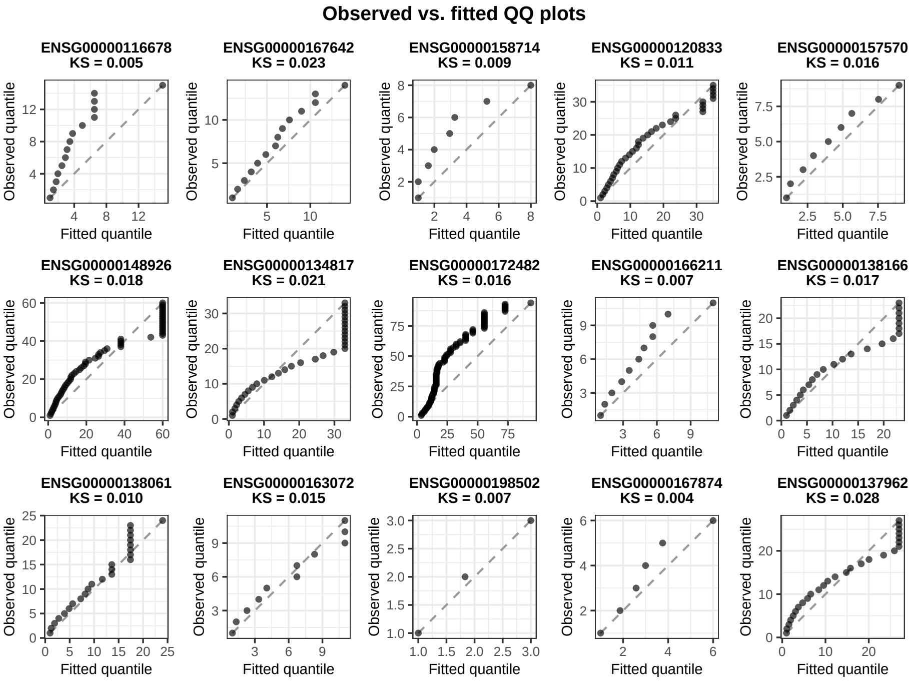

**Figure S2: Quantile–quantile plot of the NBMM fit on the IFALD dataset.**The plot compares empirical quantiles of the observed expression distribution with quantiles implied by the fitted NBMM. Closer agreement along the diagonal indicates better calibration of the fitted expression model.

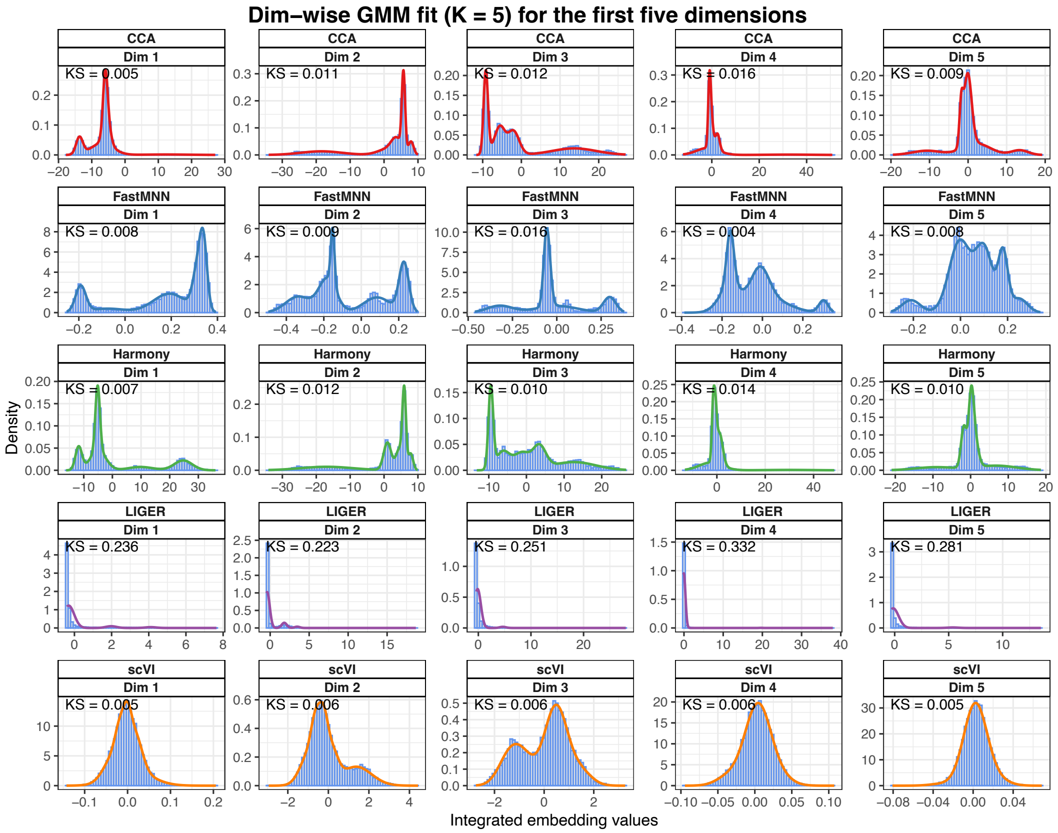

**Figure S3: Dimension-wise Gaussian mixture model fits for integrated embeddings on the IFALD dataset.** For each integration method, the distribution of embedding values is shown for the first five dimensions. Blue histograms represent the empirical distribution of integrated embedding values, and the overlaid colored density curves represent the fitted Gaussian mixture model with $K = 5$ components. Each panel reports the corresponding integration method, embedding dimension, and Kolmogorov–Smirnov (KS) statistic between the observed and fitted distributions. Smaller KS values indicate better agreement between the fitted mixture model and the empirical embedding distribution.

#### 2.2 Empirical Validation on MERFISH Data

To further assess whether the distributional assumptions used in IntegrateRigor extend to spatial transcriptomics data, we performed the same empirical validation on the Allen mouse brain MERFISH dataset. For the NBMM component, we compared the observed nonzero gene-expression count distributions with the fitted distributions and summarized the discrepancy using the KS statistic. The resulting QQ plots and KS statistics show that the NBMM provides a reasonable approximation to the empirical expression distributions for most genes, supporting its use for modeling spatial transcriptomic counts.

We next evaluated the GMM assumption for the integrated embedding. As in the single-cell analysis, we fitted Gaussian mixture models to embedding dimensions and compared the fitted densities with the empirical distributions. The density plots and KS statistics indicate generally good agreement between the fitted GMMs and the observed embedding distributions, except for the LIGER method. Together, these results suggest that the core distributional assumptions of IntegrateRigor remain empirically reasonable in the MERFISH spatial transcriptomics setting.

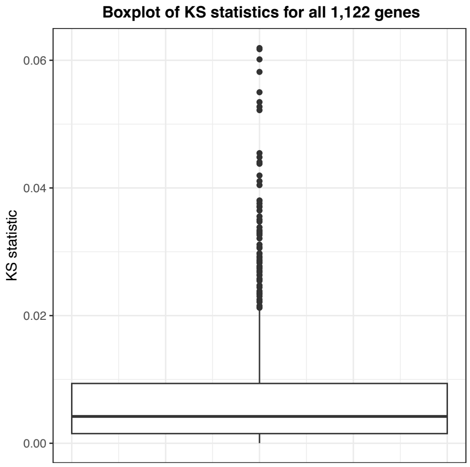

(a) NBMM KS statistics

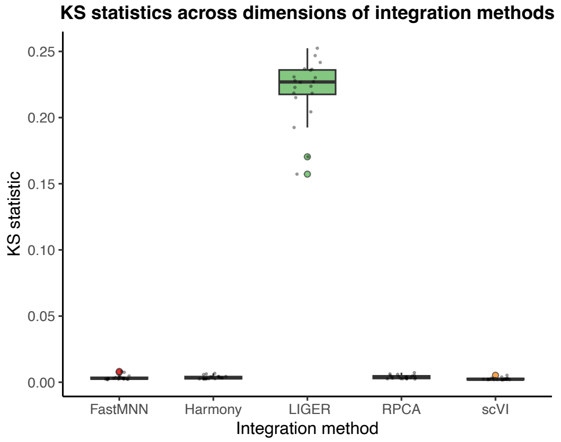

(b) GMM KS statistics

**Figure S4: Goodness-of-fit assessment using KS statistics on the Allen mouse brain MERFISH dataset.** Left panel: Distribution of KS statistics for all 1,122 genes fitted by the negative binomial mixture model (NBMM), showing generally small distributional discrepancies between the observed and fitted gene expression distributions. Right panel: Distribution of KS statistics across embedding dimensions for Gaussian mixture model (GMM) fits to integrated embeddings from different integration methods. Most methods show small KS statistics, whereas LIGER exhibits substantially larger KS values, indicating poorer compatibility with the GMM assumption.

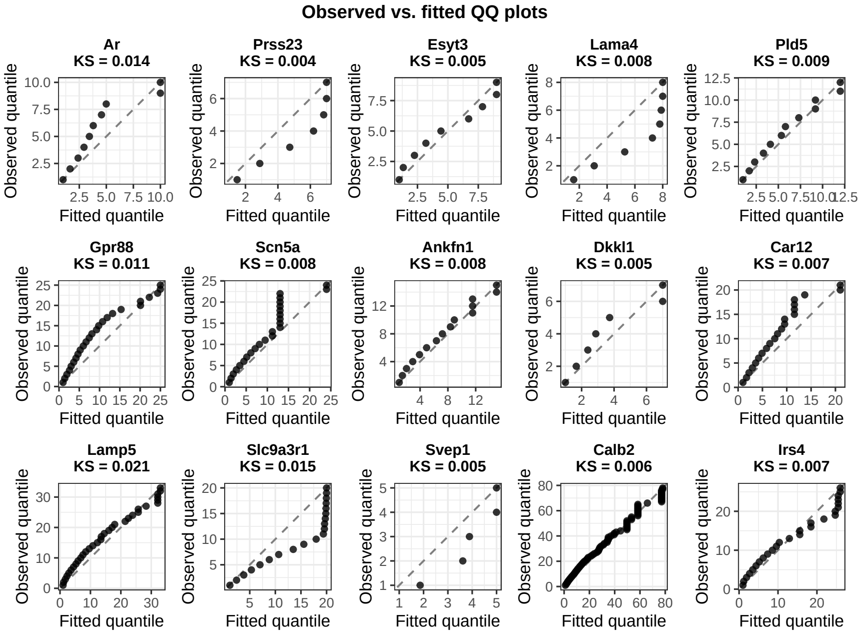

**Figure S5: Quantile–quantile plot of the NBMM fit on the Allen mouse brain MERFISH dataset.** The plot compares empirical quantiles of the observed expression distribution with quantiles implied by the fitted NBMM. Closer agreement along the diagonal indicates better calibration of the fitted expression model.

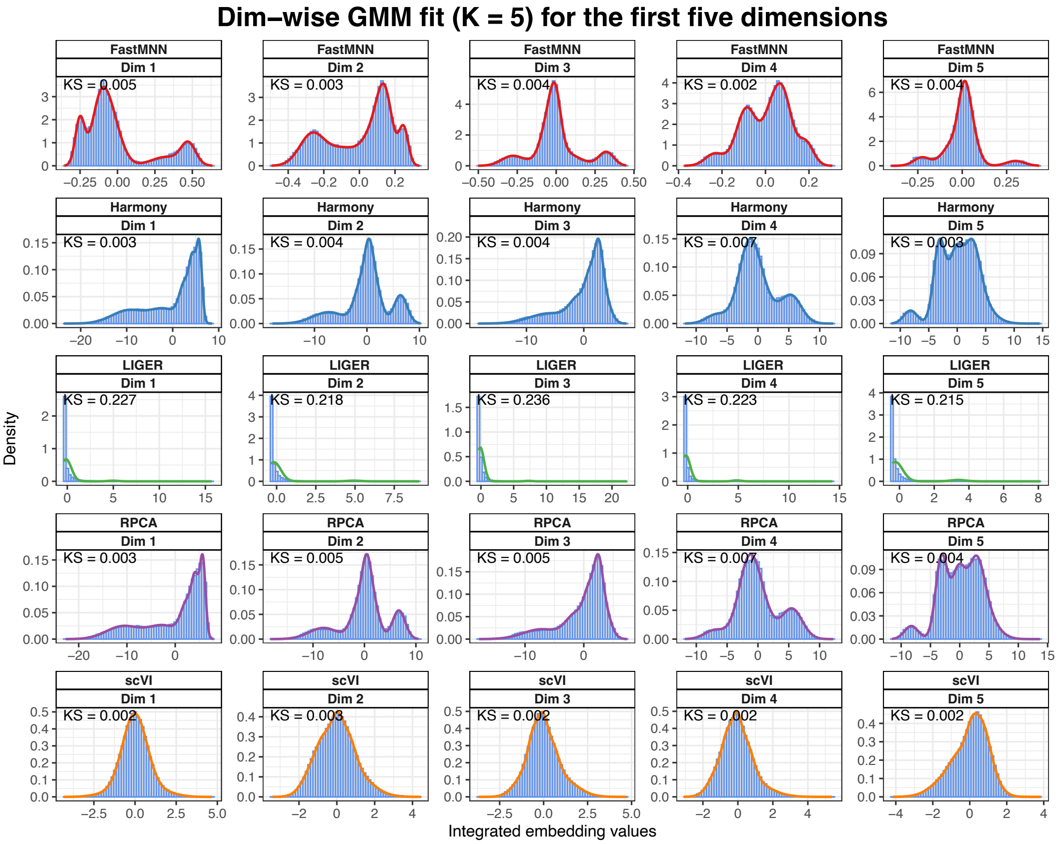

**Figure S6: Dimension-wise Gaussian mixture model fits for integrated embeddings on the Allen mouse brain MERFISH dataset.** For each integration method, the distribution of embedding values is shown for the first five dimensions. Blue histograms represent the empirical distribution of integrated embedding values, and the overlaid colored density curves represent the fitted Gaussian mixture model with K = 5 components. Each panel reports the corresponding integration method, embedding dimension, and Kolmogorov–Smirnov (KS) statistic between the observed and fitted distributions. Smaller KS values indicate better agreement between the fitted mixture model and the empirical embedding distribution.

### 3 Pearson correlation between different integrated embedding dimensions

Since the embedding-level integration score is obtained by summing dimension-wise scores, we examined whether different embedding dimensions are approximately orthogonal in practice. For each integration method, we calculated the absolute Pearson correlation between all pairs of integrated embedding dimensions.

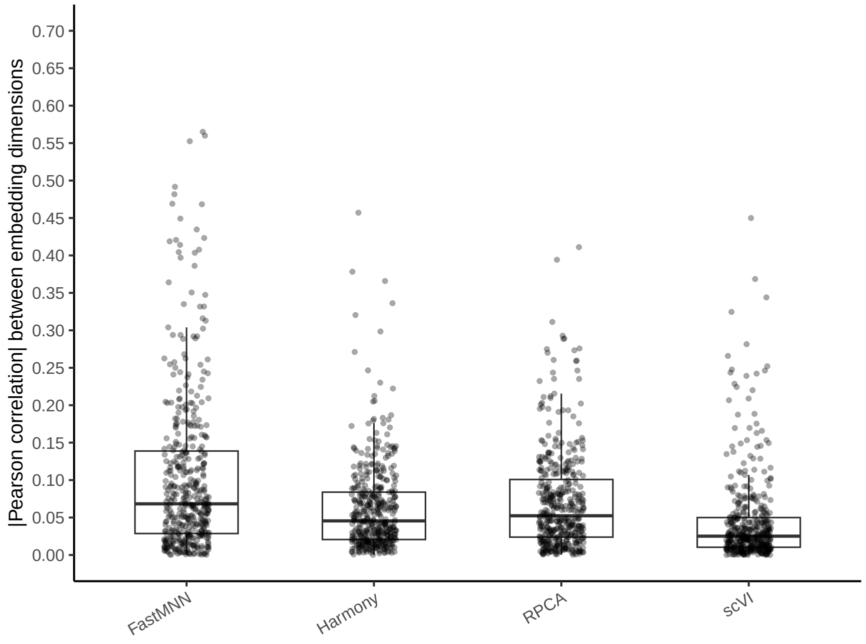

**Figure S7: Pairwise Pearson correlations between integrated embedding dimensions.** For each integration method, we calculated the absolute Pearson correlation between all pairs of embedding dimensions. Each point represents one pair of dimensions, and boxplots summarize the distribution across all dimension pairs.

### 4 Reference-batch choice stability

IntegrateRigor requires selecting a reference batch that contains all cell identities. In some datasets, however, multiple batches may satisfy this requirement and therefore can serve as candidate reference batches. To evaluate the sensitivity of IntegrateRigor to the choice of reference batch, we used the IFALD dataset, in which all three batches contain the major cell identities (C54 missing the dendritic cells) and can therefore be considered potential references. The cell numbers for each cell-identity within each batch are summarized in Table S2. By default, the IntegrateRigor implementation automatically selected IFALDA073 as the reference batch. We then manually repeated the analysis using the other candidate batches as references to assess the stability of the selected batch-stable genes and downstream integration results. We showed the following results: (1) the Jaccard similarity between the BSGs when different batches are selected as reference; (2) the integration score vs. different hyper-parameter values for Harmony when different batches are selected as reference.

**Table S2:** Cell numbers for each cell identity across IFALD batches. Rows correspond to batches, and columns correspond to annotated cell identities.

| Batch | HSC | T | Kupffer | Mono phagocyte | Hepatocyte | Erythrocyte | Platelet | Macrophage | NK | Mature B | Plasma | $\gamma\delta$ T | cDC | LSEC | Cholangiocyte | Cycl. myeloid | Cycl. plasma |
|---|---|---|---|---|---|---|---|---|---|---|---|---|---|---|---|---|---|
| C54 | 124 | 354 | 279 | 60 | 2532 | 3 | 2 | 74 | 78 | 68 | 34 | 151 | 0 | 502 | 20 | 9 | 5 |
| C96 | 302 | 1043 | 418 | 800 | 58 | 1 | 6 | 484 | 532 | 112 | 21 | 700 | 20 | 1338 | 6 | 19 | 0 |
| IFALD073 | 95 | 945 | 534 | 564 | 112 | 0 | 4 | 208 | 1100 | 893 | 21 | 363 | 33 | 772 | 37 | 54 | 5 |

**Table S3: The Jaccard similarity between the BSGs when different batches are selected as reference.**

| | C96 | IFALD073 | C54 |
|---|---|---|---|
| C96 | 1.000 | 0.987 | 0.986 |
| IFALD073 | 0.987 | 1.000 | 0.992 |
| C54 | 0.986 | 0.992 | 1.000 |

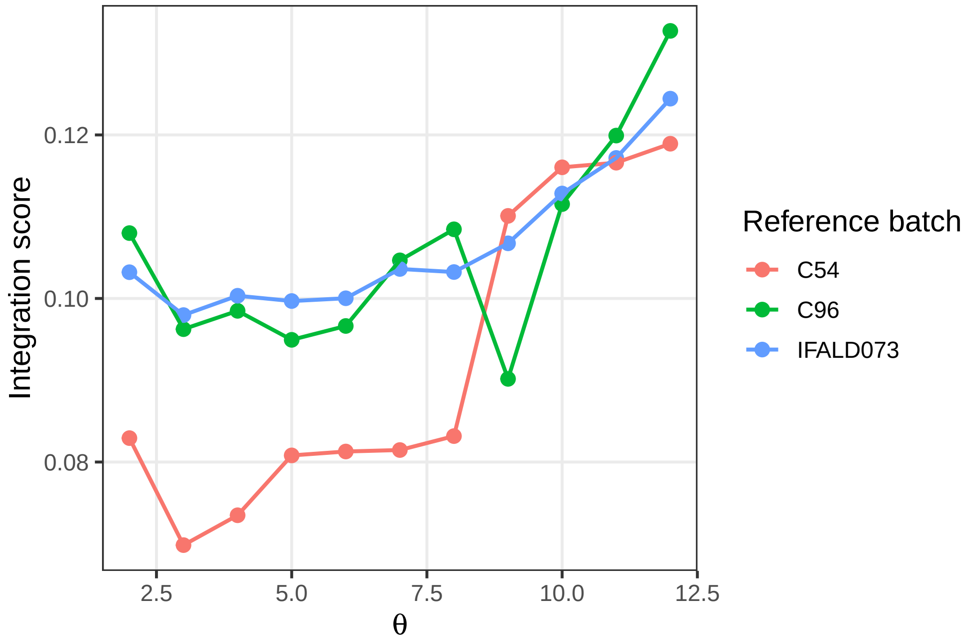

**Figure S8: The integration score versus different hyper-parameter ($\theta$) values for Harmony when different batches are selected as reference.**

### 5 Genes contributing to batch effect in the unintegrated PCA overlaps with the batch-unstable genes

We examined whether genes identified by IntegrateRigor as batch-unstable correspond to genes that visually drive batch separation before integration. To do this, we generated a PCA biplot using the unintegrated gene expression matrix. In this plot, each point represents a cell projected onto the first two principal components, with colors indicating batch identity. The arrows represent the top 100 genes with the largest contributions to the PCA axes; the direction and length of each arrow indicate how strongly and in which direction each gene contributes to variation in the PCA space.

The biplot shows clear batch-associated structure in the unintegrated embedding. Importantly, many genes whose loading directions align with batch-separated regions are classified by IntegrateRigor as batch-unstable. In contrast, batch-stable genes are less concentrated along the major batch-separating directions. This overlap supports the interpretation that IntegrateRigor identifies genes whose variation is driven by batch effects and should therefore be down-weighted or excluded during integration-specific feature selection.

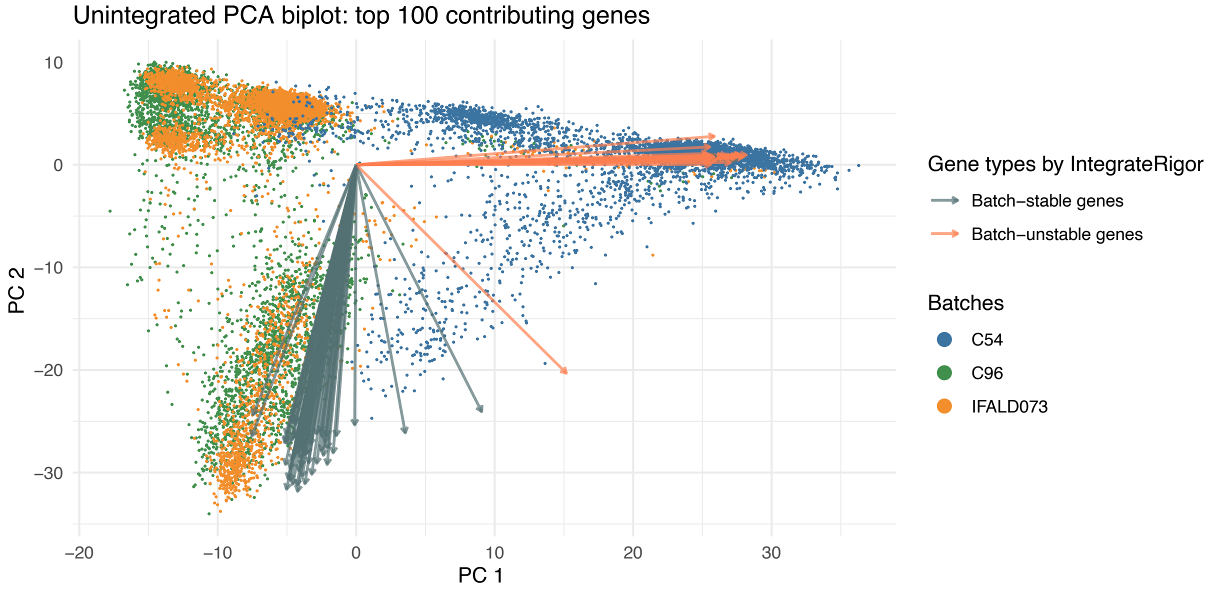

**Figure S9: PCA biplot for batch-stability assessment of top contributing genes in the unintegrated PCA space.** The unintegrated PCA embedding is shown for cells from three batches, with cells colored by batch. Arrows represent the top 100 genes contributing to the PCA structure, and arrow colors indicate whether each gene is classified by IntegrateRigor as batch-stable or batch-unstable. Batch-unstable genes tend to align with directions separating batches, whereas batch-stable genes are less associated with batch-driven variation, supporting the use of IntegrateRigor for identifying genes that are less affected by batch effects before integration.

### 6 Batch stability score correlates with annotation-dependent GTE scores

To validate the gene-level batch stability score used in IntegrateRigor, we also compared it with an existing annotation-dependent measure of gene-level technical effects (GTEs). GTEs quantify the extent to which each gene is affected by technical variation across batches or technologies, but their estimation relies on prior cell identity annotations. In contrast, IntegrateRigor estimates gene-wise batch stability directly from the observed expression distribution without requiring cell type or state labels. Therefore, a strong positive association between IntegrateRigor's batch-instability score and GTEs would support the validity of our annotation-free scoring strategy.

Across six datasets, we observed consistently positive correlations between the negative batch stability score from IntegrateRigor and the overall technical effects estimated by GTEs. Each point represents one gene, and both quantities are shown on the log scale. IntegrateRigor tends to assign higher batch-instability scores to genes with stronger annotation-dependent GTE scores. The Pearson correlations ranged from moderate to strong across datasets and were statistically significant in all cases.

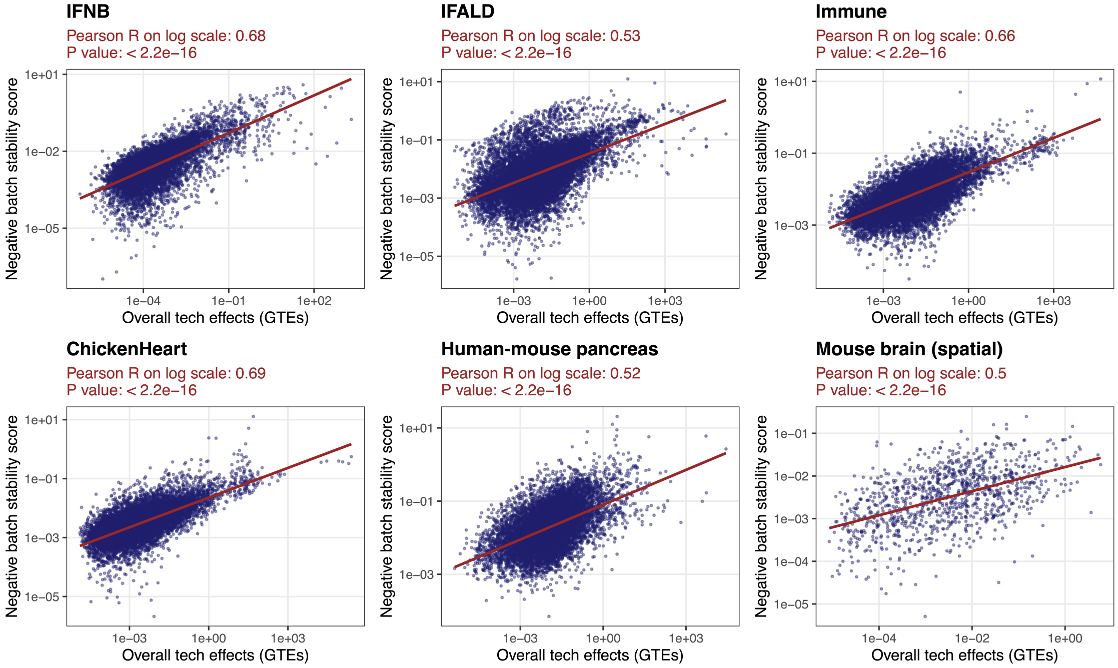

**Figure S10: Correlation between gene-wise technical effects and IntegrateRigor batch-instability scores across datasets.** Each panel shows the relationship between the overall gene-level technical effect estimated by GTEs and the negative batch stability score computed by IntegrateRigor on the log scale. Each point represents one gene, and both axes are shown on the log scale. The red line indicates the fitted linear trend. Across all six datasets, genes with larger estimated technical effects tend to have higher negative batch stability scores, with significant positive Pearson correlations. This agreement suggests that IntegrateRigor effectively identifies genes whose expression variation is strongly associated with batch- or technology-driven effects.

### 7 IntegrateRigor identifies ribosomal genes more frequently as batch-unstable genes

We examined the biological and technical characteristics of genes classified by IntegrateRigor as batch-stable genes (BSGs) or batch-unstable genes (BUSGs). Because ribosomal and mitochondrial genes are commonly associated with technical variation, sequencing depth, cell quality, or sample-processing differences, we evaluated whether these gene categories were enriched among BUSGs across datasets.

We notice that ribosomal genes were consistently enriched among batch-unstable genes across datasets(Table S4), but individual ribosomal genes differed in their stability. For the four datasets, i.e., datasets excluding the cross-species and spatial datasets, there were still 15 ribosomal genes repeatedly classified as batch-stable (Fig. S11). To better understand this difference in stability among ribosomal genes, we compared ribosomal genes identified as unstable in at least one dataset with ribosomal genes that remained stable across all datasets. The unstable ribosomal genes (e.g. *RPL21*, *RPL3*, *RPL5*, *RPL26*, *RPS3*, *RPS6*) were genes encoding the core structural components of the ribosome, whereas the consistently stable ribosomal genes (e.g. *RPS27L*, *RPL22L1*, *RPS6KB1*, *RPS6KB2*) were enriched for ribosome-related genes with additional or specialized functions, such as ribosome-associated regulation [8–11]. This suggests that ribosomal genes are not uniformly sensitive to between-batch variations, consistent with previous gene-wise analysis [12].

| | Counts | | Ribosomal genes (%) | | Mitochondrial genes (%) | |
|---|---|---|---|---|---|---|
| Dataset | BSG | BUSG | BSG | BUSG | BSG | BUSG |
| IFNB | 8831 | 14018 | 0.85 | 6.82 | 0.00 | 0.00 |
| IFALD | 10522 | 10136 | 0.34 | 14.50 | 0.03 | 2.72 |
| Immune | 264 | 331 | 0.59 | 29.79 | 0.00 | 0.00 |
| ChickenHeart | 94 | 78 | 0.56 | 29.49 | 0.08 | 5.13 |

**Table S4: Ribosomal and mitochondrial gene enrichment among IntegrateRigor-defined batch-stable and batch-unstable genes across datasets.** Here, BSG denotes batch-stable genes. BUSG denotes batch-unstable genes.

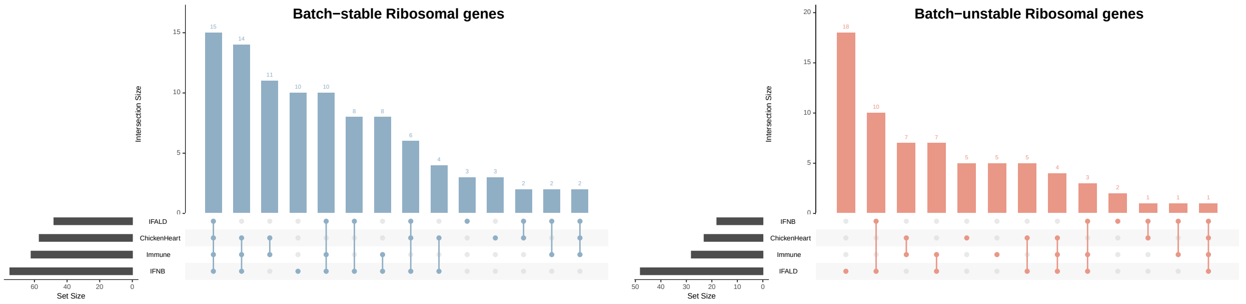

**Figure S11: Cross-dataset overlap of ribosomal genes classified as batch-stable or batch-unstable by IntegrateRigor.** Upset plots showing the intersection patterns of ribosomal genes identified as batch-stable genes (BSGs; top or batch-unstable genes (BUSGs; bottom) across datasets. The horizontal bars indicate the total number of ribosomal genes in each dataset-specific BSG or BUSG set, while the vertical bars indicate the size of each cross-dataset intersection. Connected dots below each vertical bar denote the datasets included in the corresponding intersection. The results show that ribosomal genes classified as BUSGs are repeatedly shared across datasets, supporting their association with batch-driven technical variation.

#### 8 Metrics evaluation for different stages of IntegrateRigor

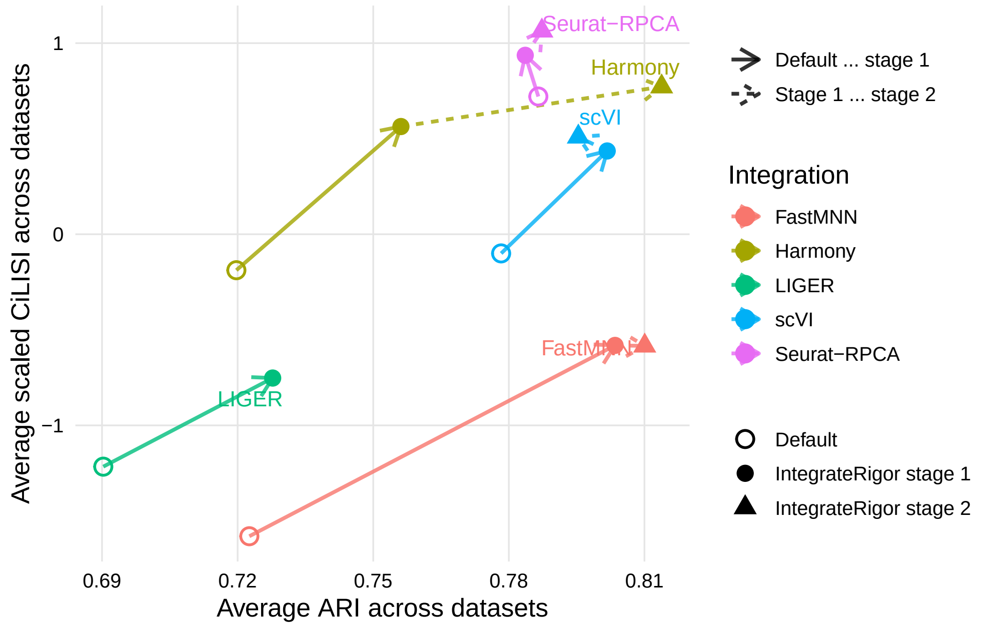

**Figure S12: Average integration performance across datasets, comparing default integration with IntegrateRigor after stage 1 and after stage 2.**

#### 9 Figures with complete legends for main figures

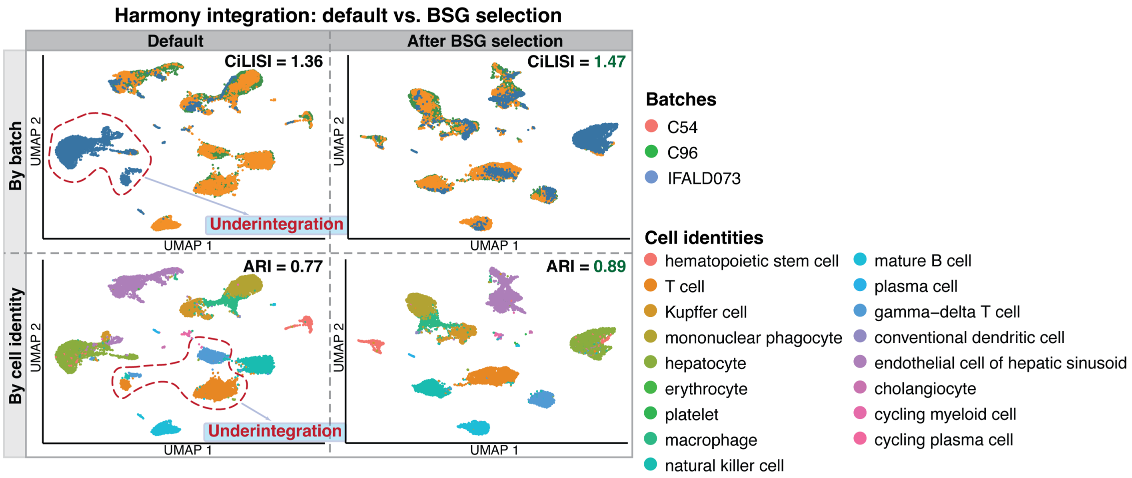

**Figure S13: Main Figure 2b with complete batch and cell identity labels**

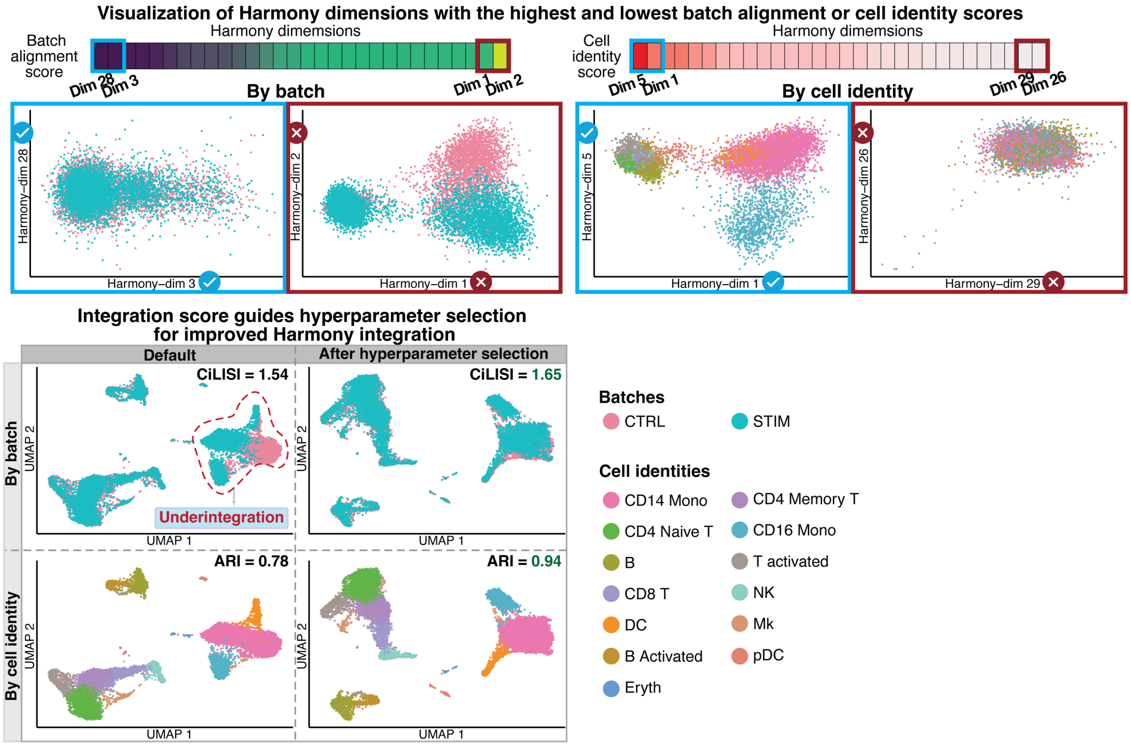

**Figure S14: Main Figure 2 f-g with complete batch and cell identity labels**

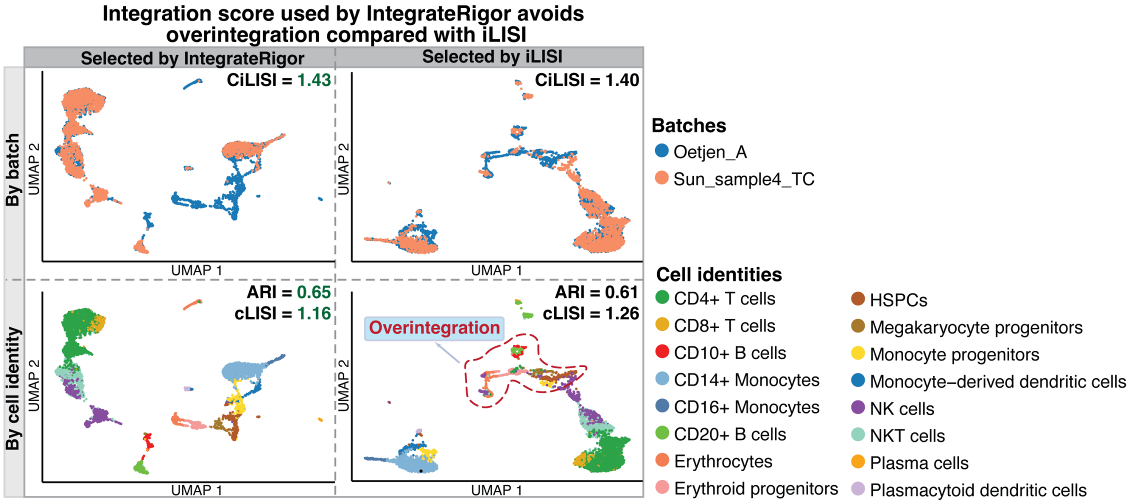

**Figure S15: Main Figure 2h with complete batch and cell identity labels**

### 10 Harmony parameters selected by existing metrics

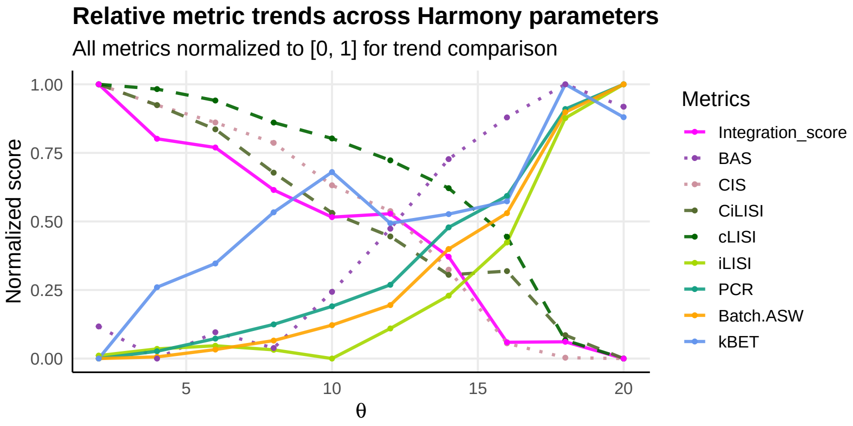

**Figure S16: Relative metric trends across Harmony integration parameters.** Line plot showing how integration performance metrics change across different values of the Harmony parameter $\theta$. In Harmony, it controls the strength of the diversity clustering penalty, with larger values encouraging stronger batch mixing during integration. All metrics are min–max normalized to the range $[0, 1]$ to facilitate comparison of relative trends across metrics. The IntegrateRigor integration score peaks at moderate $\theta$ values and decreases as $\theta$ becomes larger, suggesting that overly strong batch correction may reduce biological structure preservation. In contrast, several conventional batch-mixing metrics, including LISI, PCR, ASW, and kBET, continue to improve with larger $\theta$, highlighting their tendency to favor stronger integration even when it may lead to over-integration.

### 11 IntegratRigor sensitivity and convergence to the IntegrateRigor hyperparameters

We evaluated the stability of IntegrateRigor outputs across key model-fitting settings, including the number of mixture components and random initialization seeds, and assessed convergence across different maximum numbers of EM iterations. For each setting, we compared the resulting gene-wise batch stability scores (BSSs) and selected batch-stable genes (BSGs) with those obtained under the default configuration.

Across different choices of the number of mixture components, BSG selection remained highly consistent with the default result. The UpSet plot shows that most BSGs were shared across the parameter K, the number of mixture components, and the Jaccard similarity between each BSG set, and the default BSG set remained close to 1. Similarly, the gene-wise BSSs were highly correlated with those from the default K, indicating that the relative gene stability scores are stable across different mixture model specifications. We next examined convergence with respect to the number of EM algorithm's iterations per gene. As the iteration limit increased, both the selected BSG sets and gene-wise BSSs remained relatively unchanged compared to the default setting. The high Jaccard similarity of BSGs and high Pearson correlation of BSSs indicate that the estimates had already relatively converged under the default iteration setting, and further increasing the number of iterations produced minimal changes.

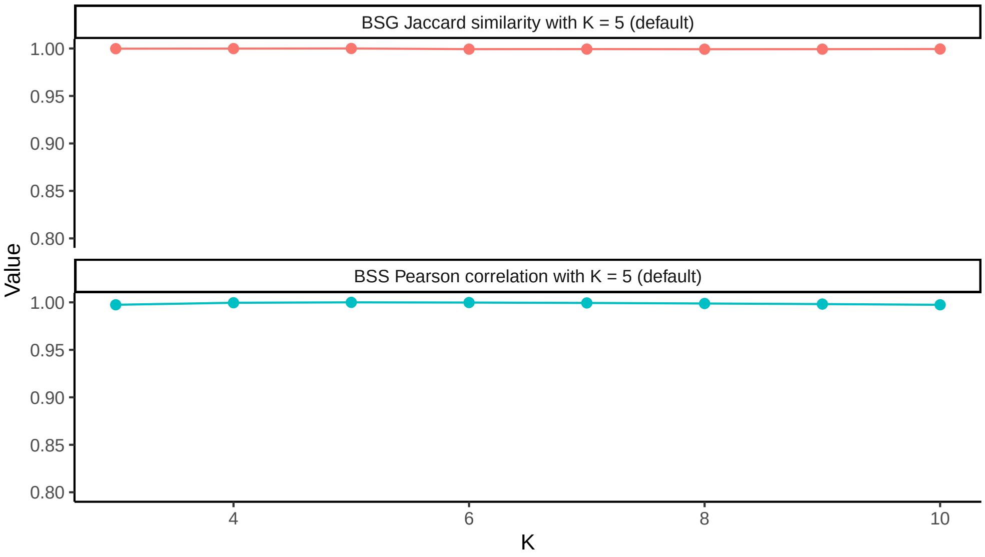

**Figure S17: Stability of BSG selection and batch stability scores across different numbers of mixture components.** Line plots showing the robustness of IntegrateRigor outputs across different choices of K, where K = 5 is the default value. The top panel shows the Jaccard overlap between the BSGs selected at each K and those selected at K = 5. The bottom panel shows the correlation between gene-wise batch stability scores (BSSs) computed at each K and those computed at K = 5.

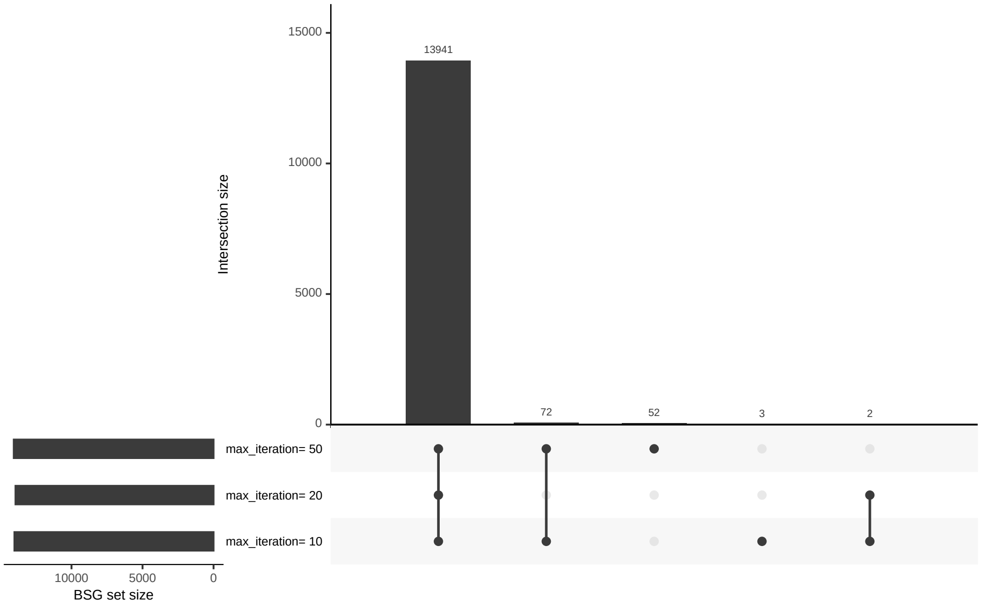

**Figure S18: Overlap of selected batch-stable genes across different numbers of EM algorithms' iterations per gene.** UpSet plot showing the overlap of batch-stable genes (BSGs) selected by IntegrateRigor under different maximum numbers of EM algorithms' iterations per gene. Horizontal bars indicate the number of BSGs selected under each iteration setting, vertical bars indicate intersection sizes, and connected dots denote the iteration settings included in each intersection.

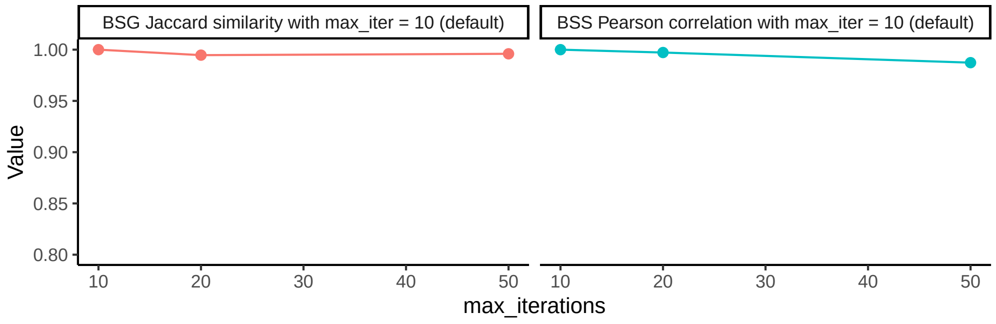

**Figure S19: Stability of BSG selection and batch stability scores across different numbers of EM algorithms' iterations per gene.** Line plots showing the convergence of IntegrateRigor outputs across different maximum numbers of EM algorithms' iterations per gene, where iterations=10 is the default value. The left panel shows the Jaccard similarity between BSGs selected at each iteration setting and those selected with the default setting. The right panel shows the Pearson correlation between gene-wise batch stability scores (BSSs) computed at each iteration setting and those computed with the default setting.

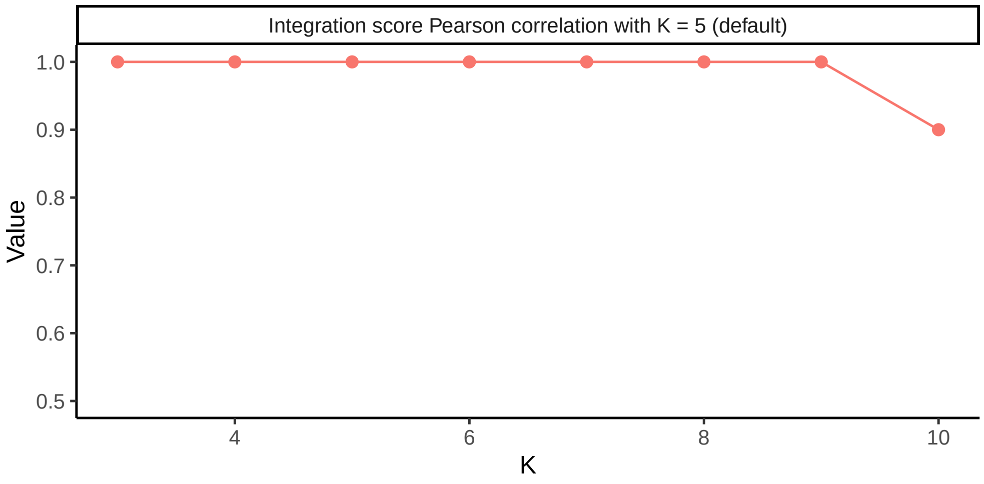

**Figure S20: Stability of integration score across different numbers of Gaussian mixture components.** Line plots showing the correlation of the integration scores across different choices of K among the same set of hyperparameter values in Harmony, where K = 5 is the default value.

### 12 Complete IntegrateRigor results

#### 12.1 Metric evaluation

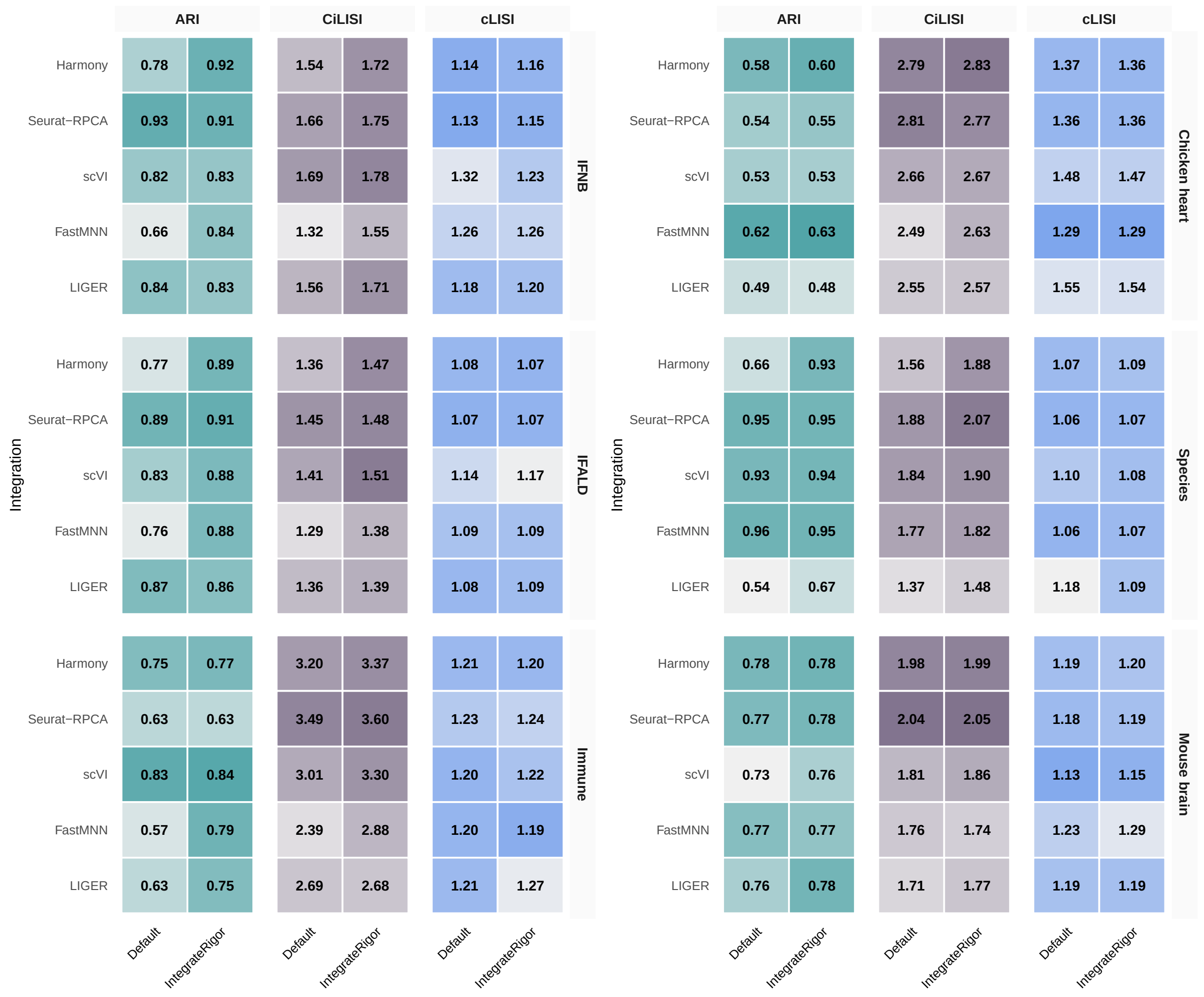

Figure S21: Metrics evaluation for IntegrateRigor vs. default integration over ARI, CiLISI, and cLISI.

#### 12.2 UMAP visualization of default integration vs. IntegrateRigor

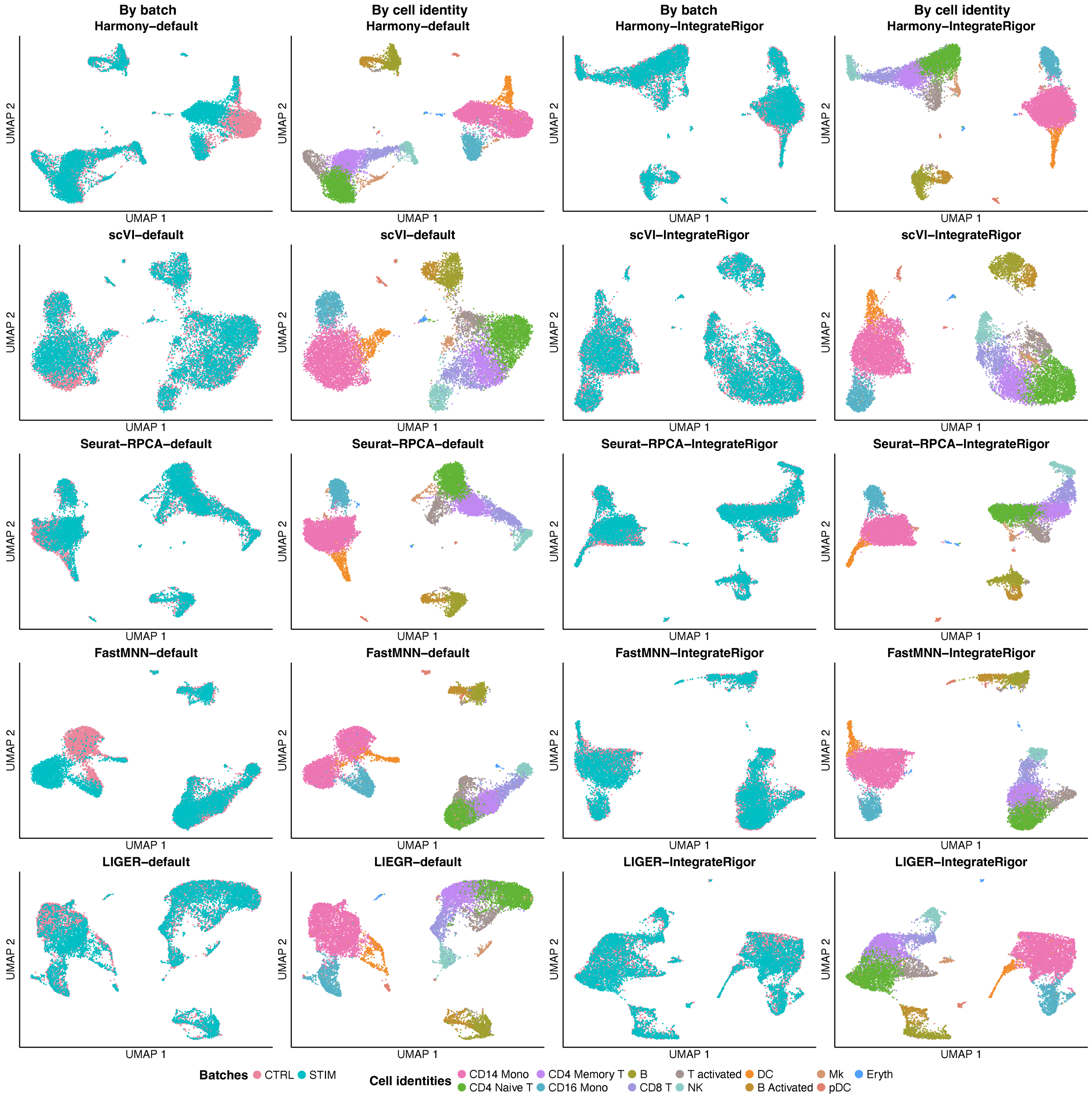

**Figure S22: Comparison of default and IntegrateRigor-selected integration results on the IFN-$\beta$ stimulated dataset.** UMAP visualizations of the dataset after integration using Harmony, scVI, Seurat-RPCA, FastMNN, and LIGER under either default feature/parameter settings or IntegrateRigor-selected settings. For each method and setting, cells/spots are shown twice: colored by batch on the left and by annotated cell type on the right.

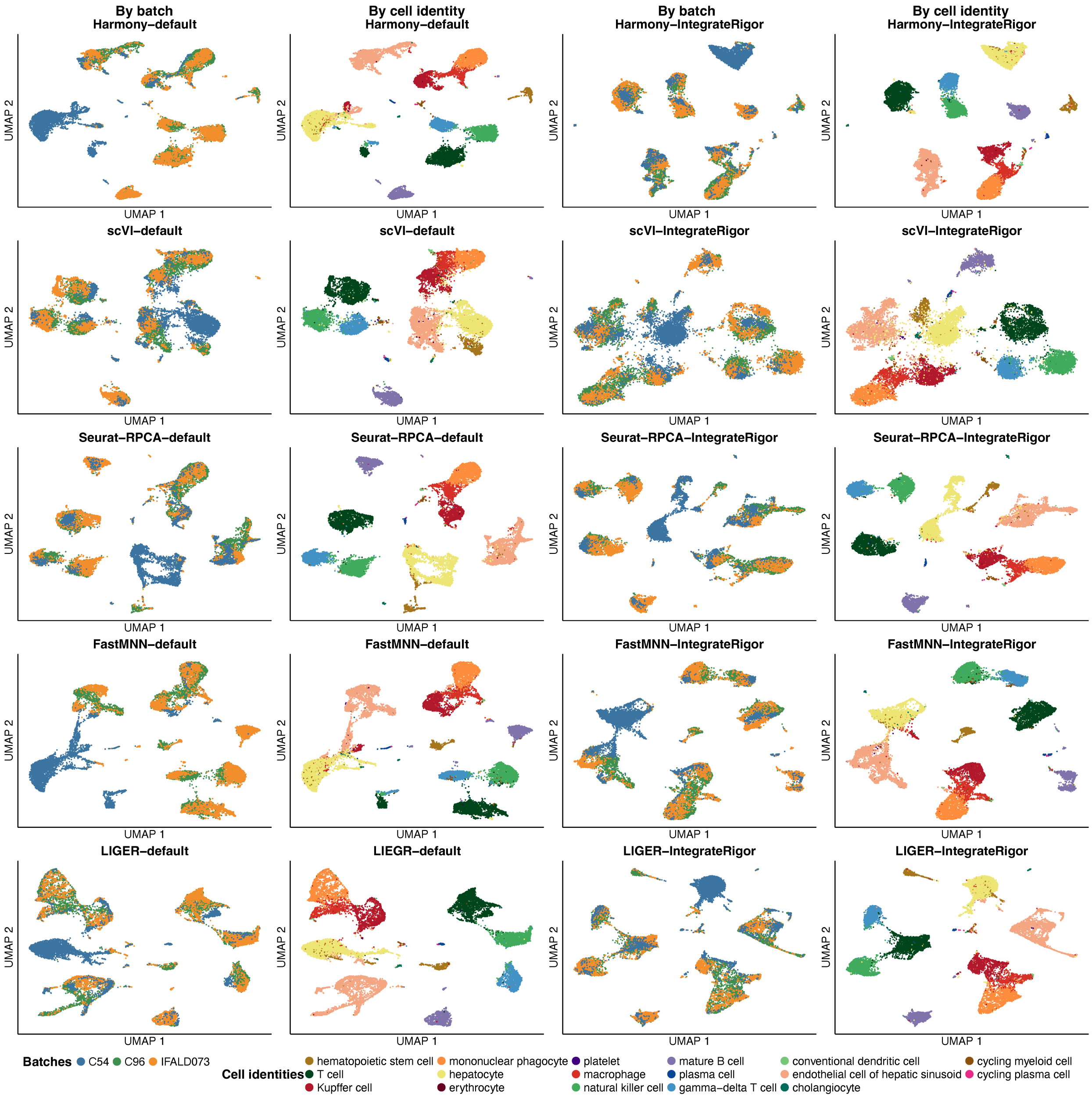

**Figure S23: Comparison of default and IntegrateRigor-selected integration results on the IFALD dataset.** UMAP visualizations of the dataset after integration using Harmony, scVI, Seurat-RPCA, FastMNN, and LIGER under either default feature/parameter settings or IntegrateRigor-selected settings. For each method and setting, cells/spots are shown twice: colored by batch on the left and by annotated cell type on the right.

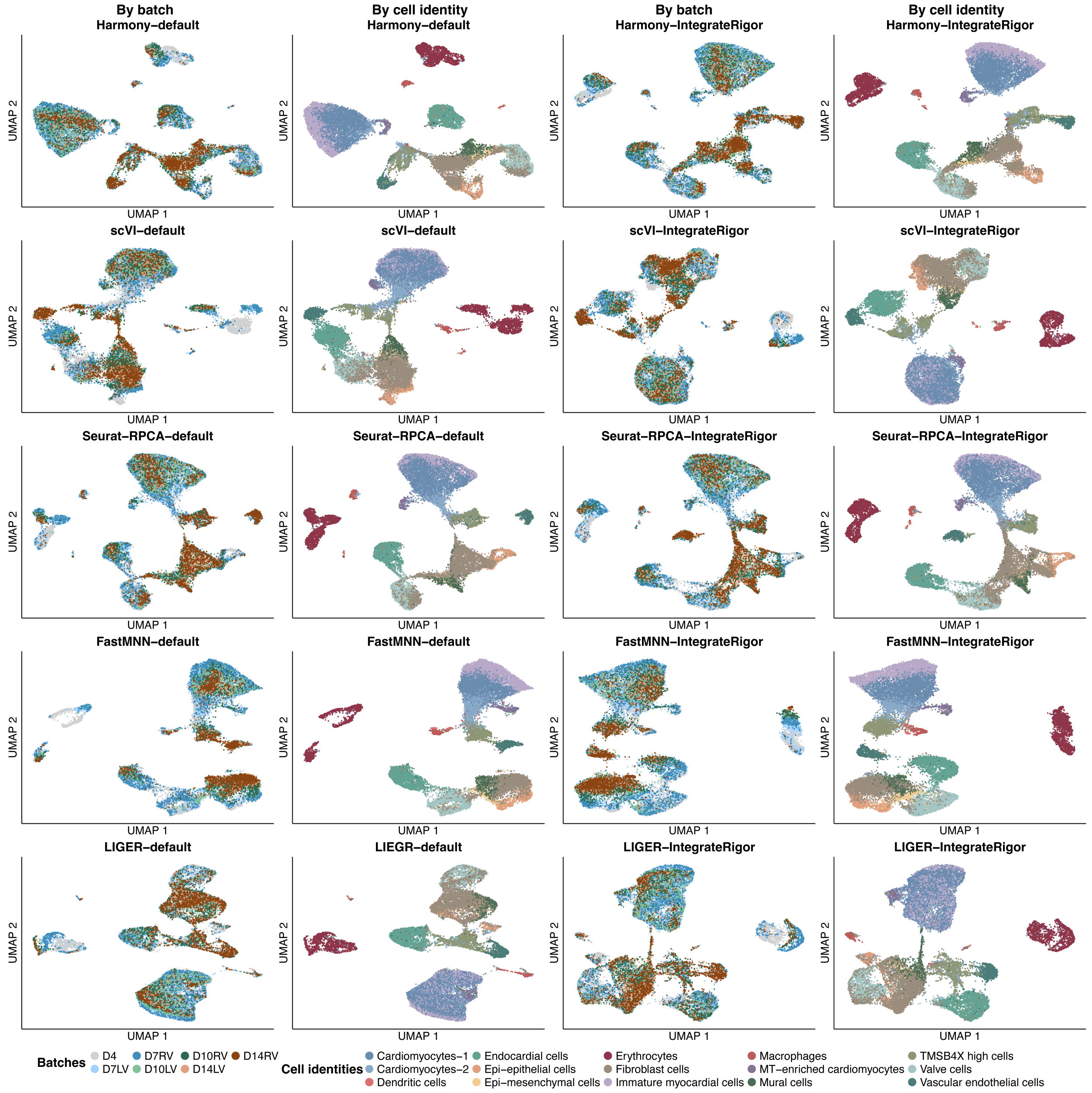

**Figure S24: Comparison of default and IntegrateRigor-selected integration results on the chicken heart dataset.** UMAP visualizations of the dataset after integration using Harmony, scVI, Seurat-RPCA, FastMNN, and LIGER under either default feature/parameter settings or IntegrateRigor-selected settings. For each method and setting, cells/spots are shown twice: colored by batch on the left and by annotated cell type on the right.

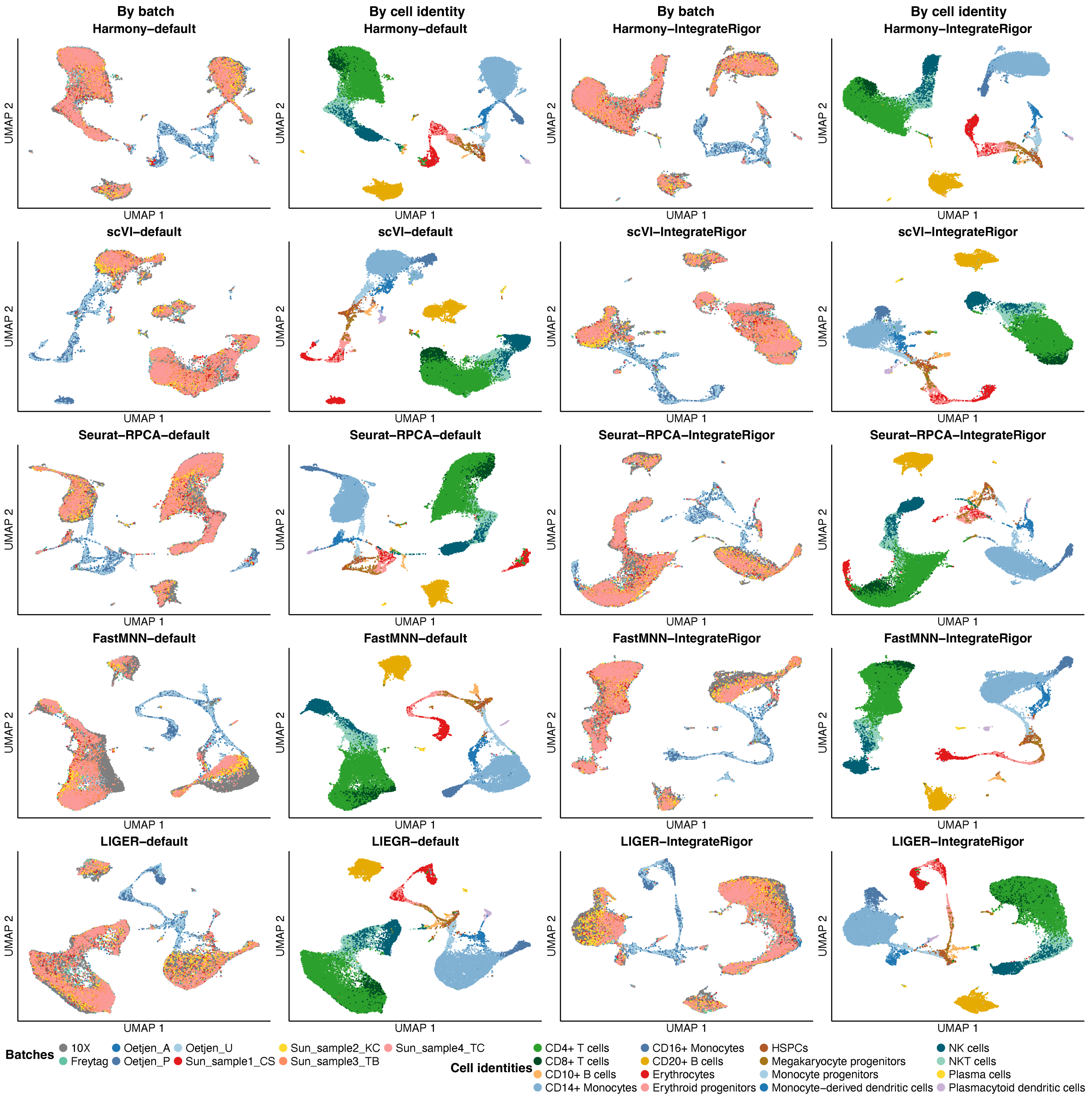

**Figure S25: Comparison of default and IntegrateRigor-selected integration results on the immune dataset.** UMAP visualizations of the dataset after integration using Harmony, scVI, Seurat-RPCA, FastMNN, and LIGER under either default feature/parameter settings or IntegrateRigor-selected settings. For each method and setting, cells/spots are shown twice: colored by batch on the left and by annotated cell type on the right.

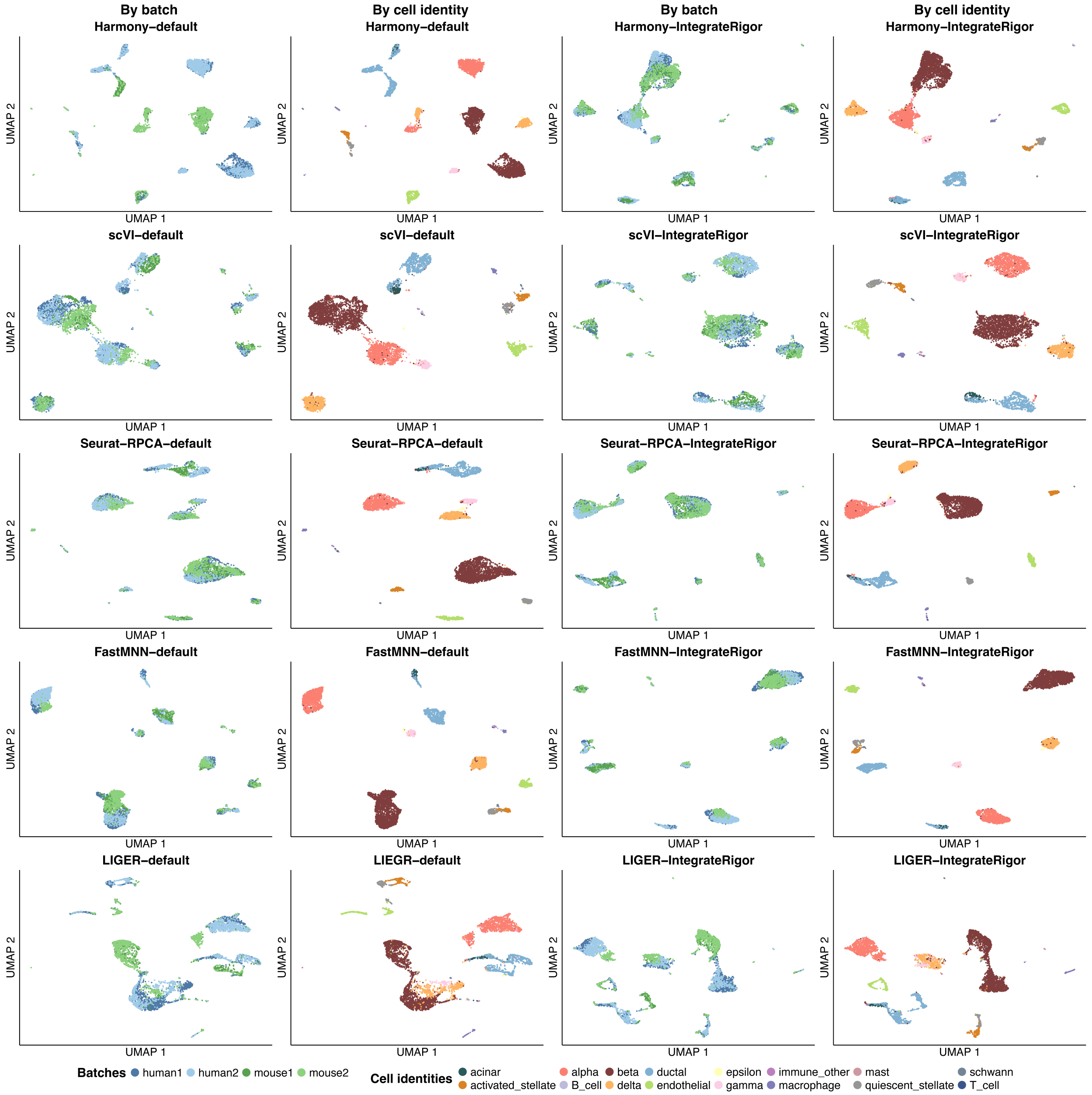

**Figure S26: Comparison of default and IntegrateRigor-selected integration results on the human-mouse pancreas dataset.** UMAP visualizations of the dataset after integration using Harmony, scVI, Seurat-RPCA, FastMNN, and LIGER under either default feature/parameter settings or IntegrateRigor-selected settings. For each method and setting, cells/spots are shown twice: colored by batch on the left and by annotated cell type on the right.

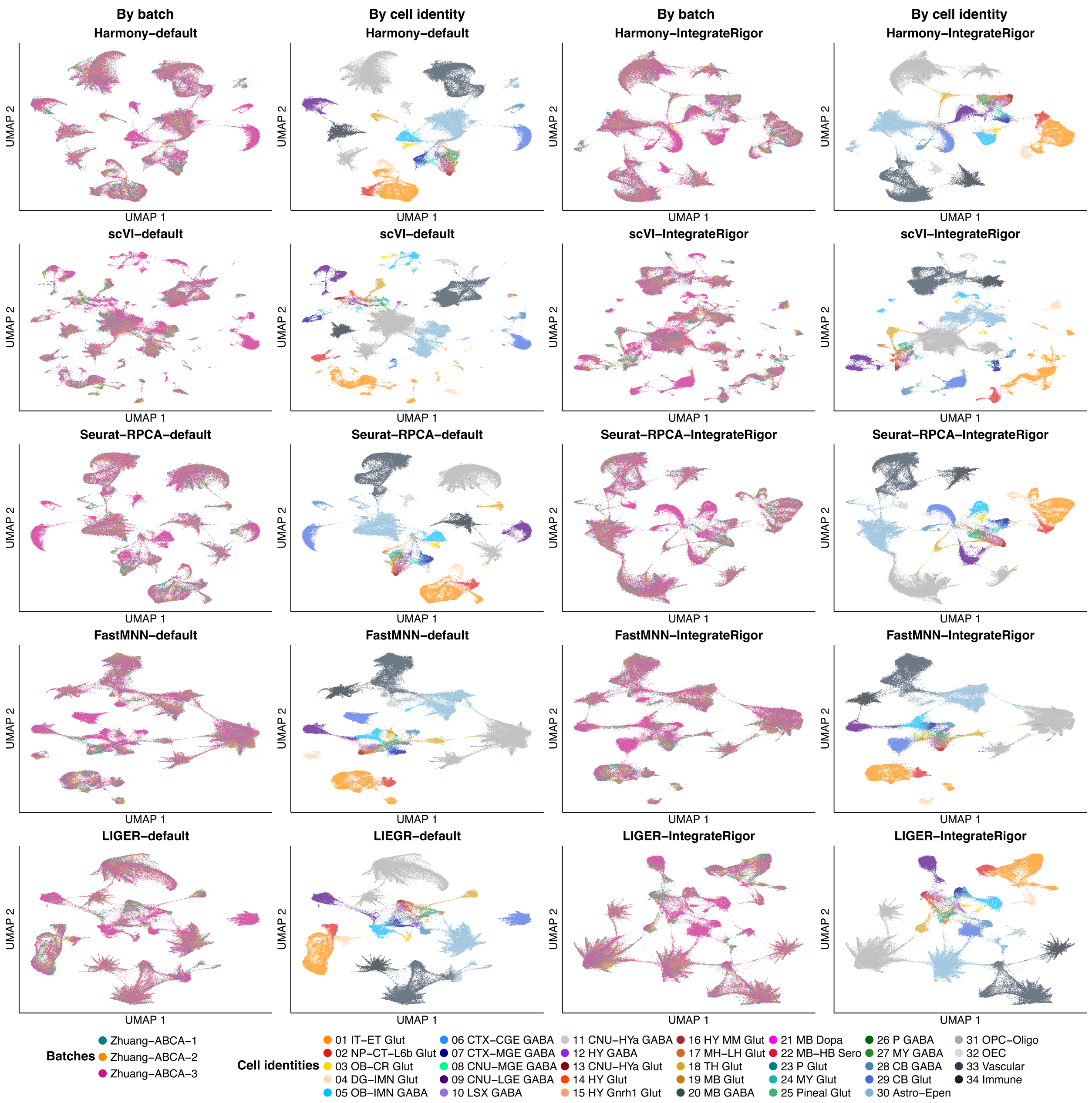

**Figure S27: Comparison of default and IntegrateRigor-selected integration results on the mouse brain spatial dataset.** UMAP visualizations of the dataset after integration using Harmony, scVI, Seurat-RPCA, FastMNN, and LIGER under either default feature/parameter settings or IntegrateRigor-selected settings. For each method and setting, cells/spots are shown twice: colored by batch on the left and by annotated cell type on the right.

### 13 Integration score across different methods

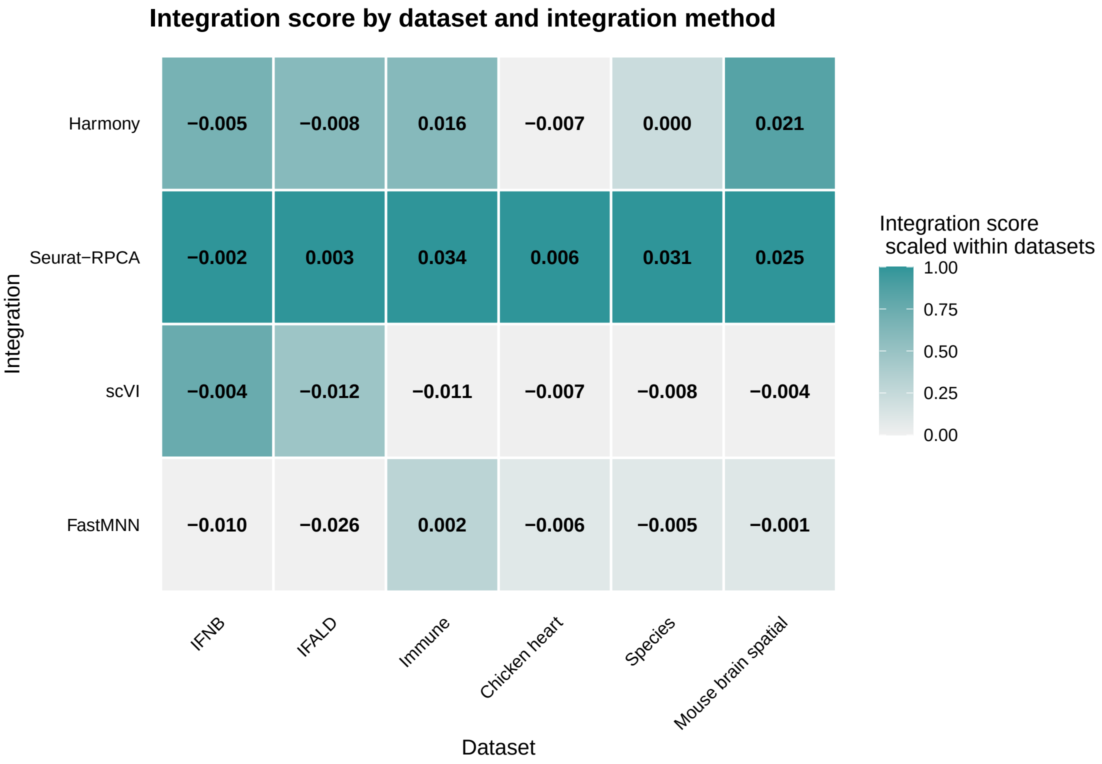

**Figure S28: Integration score values across different methods and datasets**

### 14 Cancer-immune niches in the colorectal cancer dataset

#### 14.1 Default Harmony integration results

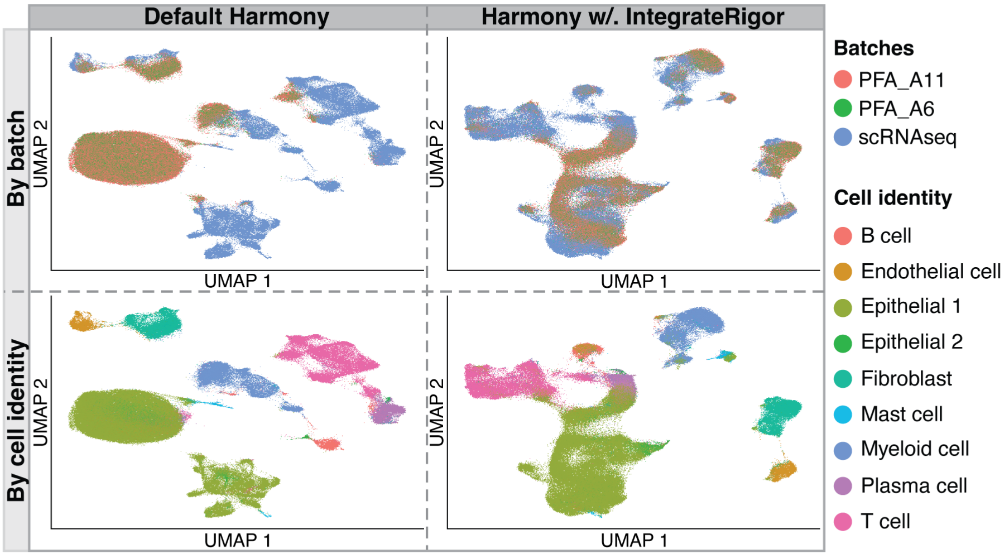

**Figure S29: Comparison of default Harmony integration and Harmony integration with IntegrateRigo for colorectal cancer dataset.** UMAP embeddings are shown for each integration strategy, with columns corresponding to default Harmony and Harmony with IntegrateRigor, and rows colored by batch or cell identity. For default Harmony, cells from different batches remain partially separated, indicating underintegration.

#### 14.2 GO terms for different immune-epithelial niches

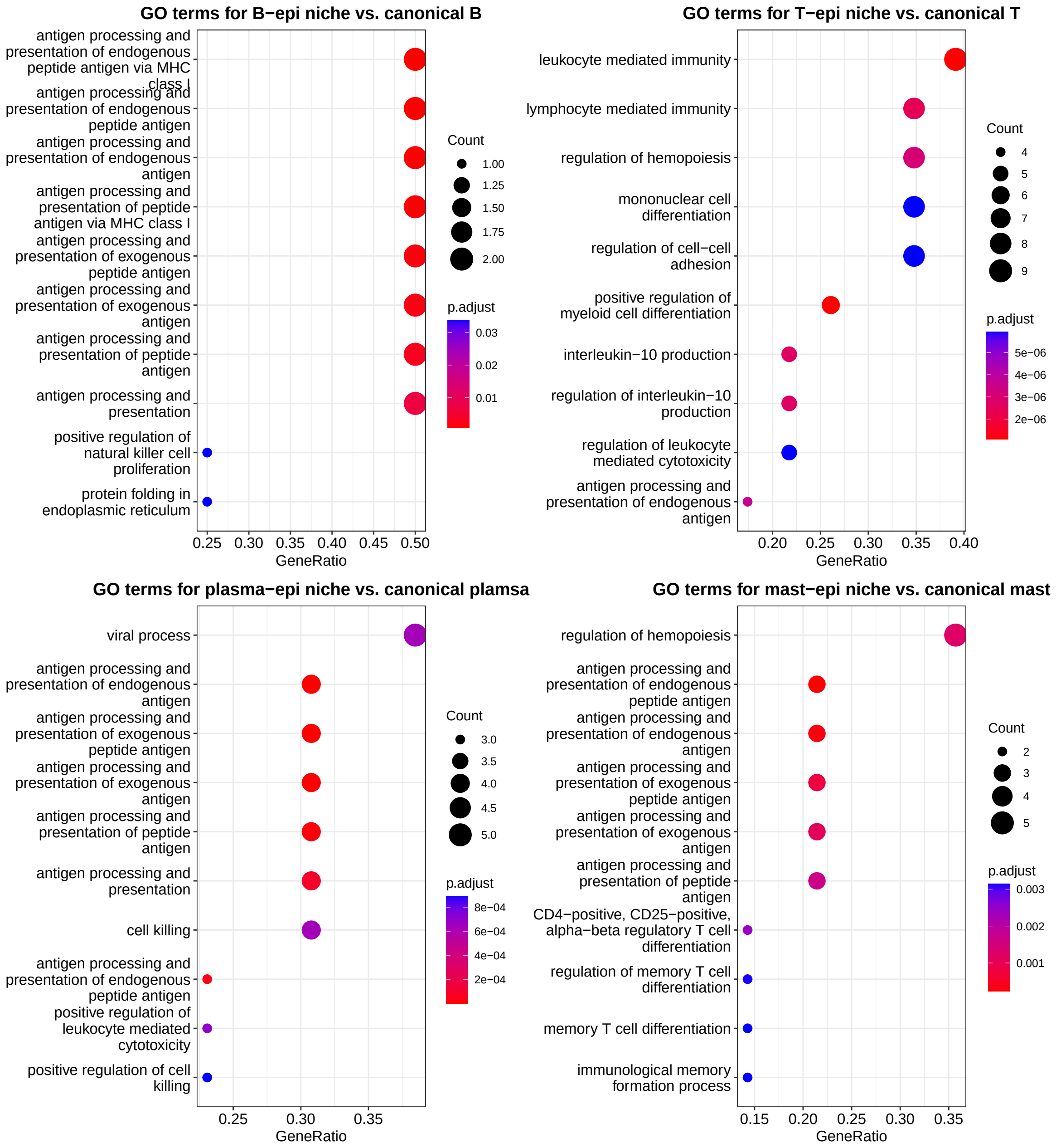

**Figure S30: GO enrichment analysis of genes distinguishing epithelial-niche cell states from canonical immune cell types.** Dot plots showing enriched Gene Ontology biological process terms for genes differentially enriched in epithelial-niche-associated B, T, plasma, and mast cell states compared with their corresponding canonical immune cell types. Each panel reports the top enriched GO terms for one comparison. The x-axis shows the gene ratio, dot size indicates the number of genes contributing to each GO term, and color represents the adjusted p-value. Across comparisons, epithelial-niche-associated cell states show enrichment for immune-related processes, including antigen processing and presentation, leukocyte-mediated immunity, lymphocyte-mediated immunity, and regulation of hematopoiesis, suggesting that these niche-associated populations retain immune functional programs while exhibiting distinct tissue-context-associated transcriptional features.
